# RNA promotes the formation of spatial compartments in the nucleus

**DOI:** 10.1101/2020.08.25.267435

**Authors:** Sofia A. Quinodoz, Prashant Bhat, Noah Ollikainen, Joanna W. Jachowicz, Abhik K. Banerjee, Peter Chovanec, Mario R. Blanco, Amy Chow, Yolanda Markaki, Kathrin Plath, Mitchell Guttman

**Affiliations:** Division of Biology and Biological Engineering, California Institute of Technology, Pasadena, CA 91125, USA; David Geffen School of Medicine, University of California, Los Angeles, Los Angeles, CA 90095, USA; Keck School of Medicine, University of Southern California, Los Angeles, CA 90089, USA; Department of Biological Chemistry, University of California, Los Angeles, Los Angeles, CA 90095, USA

**Author notes:** These authors contributed equally to this work (Prashant Bhat, Noah Ollikainen, Joanna W. Jachowicz). To whom correspondence should be addressed. (MG).

## Abstract

The nucleus is a highly organized arrangement of RNA, DNA, and protein molecules that are compartmentalized within three-dimensional (3D) structures involved in shared functional and regulatory processes. Although RNA has long been proposed to play a global role in organizing nuclear structure, exploring the role of RNA in shaping nuclear structure has remained a challenge because no existing methods can simultaneously measure RNA-RNA, RNA-DNA, and DNA-DNA contacts within 3D structures. To address this, we developed RNA & DNA SPRITE (RD-SPRITE) to comprehensively map the location of all RNAs relative to DNA and other RNAs. Using this approach, we identify many RNAs that are localized near their transcriptional loci (RNA-DNA) together with other diffusible ncRNAs (RNA-RNA) within higher-order DNA structures (DNA-DNA). These RNA-chromatin compartments span three major classes of nuclear functions: RNA processing (including ribosome biogenesis, mRNA splicing, snRNA biogenesis, and histone mRNA processing), heterochromatin assembly, and gene regulation. More generally, we identify hundreds of ncRNAs that form stable nuclear compartments in spatial proximity to their transcriptional loci. We find that dozens of nuclear compartments require RNA to guide protein regulators into these 3D structures, and focusing on several ncRNAs, we show that these ncRNAs specifically regulate heterochromatin assembly and the expression of genes contained within these compartments. Together, our results demonstrate a unique mechanism by which RNA acts to shape nuclear structure by forming high concentration territories immediately upon transcription, binding to diffusible regulators, and guiding them into spatial compartments to regulate a wide range of essential nuclear functions.

## INTRODUCTION

The nucleus is spatially organized in three-dimensional (3D) structures that are important for various functions including DNA replication, transcription, and RNA processing^1–6^. To date, genome-wide studies of nuclear organization have focused primarily on the role of DNA^2,7,8^, yet nuclear structures are known to contain multiple DNA, RNA, and protein molecules that are involved in shared functional and regulatory processes^1–6^. These include classical compartments like the nucleolus^9^ (which contains transcribed ribosomal RNAs and their processing molecules) and nuclear speckles^10^ (which contain nascent pre-mRNAs and mRNA splicing components), as well as the more recently described transcriptional condensates^11,12^ (which contain Mediator and RNA Polymerase II). Because the complete molecular architecture of the nucleus has not been globally explored, the full extent to which such nuclear compartments exist and contribute to nuclear function remains unknown. Even for the specific nuclear compartments that have been molecularly characterized, the mechanism by which RNA and protein molecules transition from diffuse locations throughout the nucleus into compartmentalized structures remains largely unknown.

Nuclear RNA has long been proposed to play a central role in shaping nuclear structure^13–18^. Initial experiments performed more than 30 years ago found that global disruption of RNA (using RNase) leads to large scale morphological deficits in the nucleus^13^. Over the past decade it has become clear that mammalian genomes encode thousands of nuclear-enriched ncRNAs^19–21^, several of which play critical roles in the regulation of essential nuclear functions^22,23^. These include ncRNAs involved in splicing of pre-mRNAs (snRNAs)^24,25^, cleavage and modification of pre-ribosomal RNAs (snoRNAs, Rnase MRP)^26–28^, 3’-end cleavage and processing of the non-polyadenylated histone pre-mRNAs (U7 snRNA)^29–32^, and transcriptional regulation (e.g. Xist^33–35^ and 7SK^36–38^). Interestingly, many of these functionally important ncRNAs localize within specific spatial compartments in the nucleus^6,39,40^. For example, snoRNAs and the 45S pre-ribosomal RNA localize within the nucleolus^9,41–43^, Xist localizes on the inactive X chromosome (Barr body)^35,44–46^, and snRNAs and Malat1 localize within nuclear speckles^10,47^.

In each of these examples, multiple RNA, DNA, and protein components simultaneously interact within precise three-dimensional structures to coordinate specific nuclear functions. While the roles of these specific ncRNAs have been well studied, comprehensively mapping the localization patterns of most nuclear ncRNAs relative to other RNAs and DNAs in 3D space remains a challenge because no existing method can simultaneously measure RNA-RNA, RNA-DNA, and DNA-DNA contacts within 3D structures. As a result, it is unclear: (i) which specific RNAs might be involved in nuclear organization^16,18,48^, (ii) which specific nuclear compartments are dependent on RNA, and (iii) what mechanisms RNA might utilize to organize nuclear structures.

Microscopy is currently the only way to relate RNA and DNA molecules in 3D space. However, this approach is limited to examining a small number of simultaneous interactions and therefore requires *a priori* knowledge of which RNAs and nuclear structures to explore. An alternative approach is genomic mapping of RNA-DNA contacts using proximity-ligation methods^49–53^. While these approaches can provide genome-wide pairwise maps of RNA-DNA interactions, they do not provide information about the 3D organization of these molecules in the nucleus. Moreover, we recently showed that proximity-ligation methods can fail to identify pairwise contacts between molecules that are organized within nuclear compartments because these methods only identify interactions where components are close enough in space to be directly ligated^54^. Consistent with this observation, existing RNA-DNA proximity-ligation methods fail to identify known RNA-DNA contacts that are contained within various well-established nuclear bodies, such as nucleoli, histone locus bodies (HLBs), and Cajal bodies^50–53^.

We recently developed SPRITE, a proximity-ligation independent method that utilizes split-and-pool barcoding to generate accurate, comprehensive, and multi-way 3D spatial maps of the nucleus across a wide range of distances^54^. Importantly, we showed that this approach can accurately map the spatial organization of DNA arranged around two nuclear bodies – nucleoli and nuclear speckles^54^. However, our original version of the technique could not detect the vast majority of ncRNAs – including low abundance ncRNAs known to organize within several well-defined nuclear structures – thereby precluding a comprehensive map of RNA localization within the nucleus. Here, we introduce a dramatically improved method, RNA & DNA SPRITE (RD-SPRITE), which enables simultaneous and high-resolution measurements of thousands of RNAs – including low abundance RNAs such as nascent pre-mRNAs and ncRNAs – relative to all other RNA and DNA molecules in 3D space. Using this approach, we identify hundreds of RNA-containing nuclear structures that are each largely organized within higher-order structures around shared regulatory targets. We demonstrate that many ncRNAs form high concentration territories within defined spatial compartments throughout the nucleus. Many of these ncRNAs bind to diffusible ncRNAs and proteins and act to guide their localization within these nuclear compartments. Focusing on several examples, we show that these ncRNAs specifically regulate genes contained within these spatial compartments. Together, our results demonstrate a privileged role for RNA in the formation of nuclear compartments that are involved in a wide range of essential nuclear functions including RNA processing, heterochromatin assembly, and gene regulation.

## RESULTS

### RD-SPRITE generates accurate maps of higher-order RNA and DNA contacts throughout the cell

Exploring the role of RNA in shaping nuclear structure has remained a challenge because no existing genomic method can simultaneously measure RNA-RNA, RNA-DNA, and DNA-DNA contacts within 3D structures. To address this, we developed RNA & DNA SPRITE (RD-SPRITE) to comprehensively map the location of all RNAs relative to DNA and other RNAs. Specifically, we improved the efficiency of the RNA-tagging steps of our SPRITE method^55^ to enable detection of all classes of RNA – from highly abundant ribosomal RNAs and snRNAs to less abundant lncRNAs and individual nascent pre-mRNAs (**Supplemental Note 1**). Briefly, our approach works as follows: (i) RNA, DNA, and protein contacts are crosslinked to preserve their spatial relationships *in situ*, (ii) cells are lysed and the contents are fragmented into smaller crosslinked complexes, (iii) DNA and RNA within each complex are tagged with a sequence-specific adaptor, (iv) barcoded using an iterative split-and-pool strategy to uniquely assign a shared barcode to all DNA and RNA components contained within a crosslinked complex, (v) DNA and RNA are sequenced, and (vi) all reads sharing identical barcodes are merged into a group that we refer to as a SPRITE cluster (**Figure 1A, Supplemental Figure 1A**, see **Methods**). Accordingly, RD-SPRITE enables simultaneous mapping of multi-way DNA-DNA, RNA-DNA, and RNA-RNA contacts in the same experiment. Because RD-SPRITE does not rely on proximity ligation, it can detect multiple RNA and DNA molecules that simultaneously associate within the nucleus (referred to as higher-order structures).

**Figure 1:**
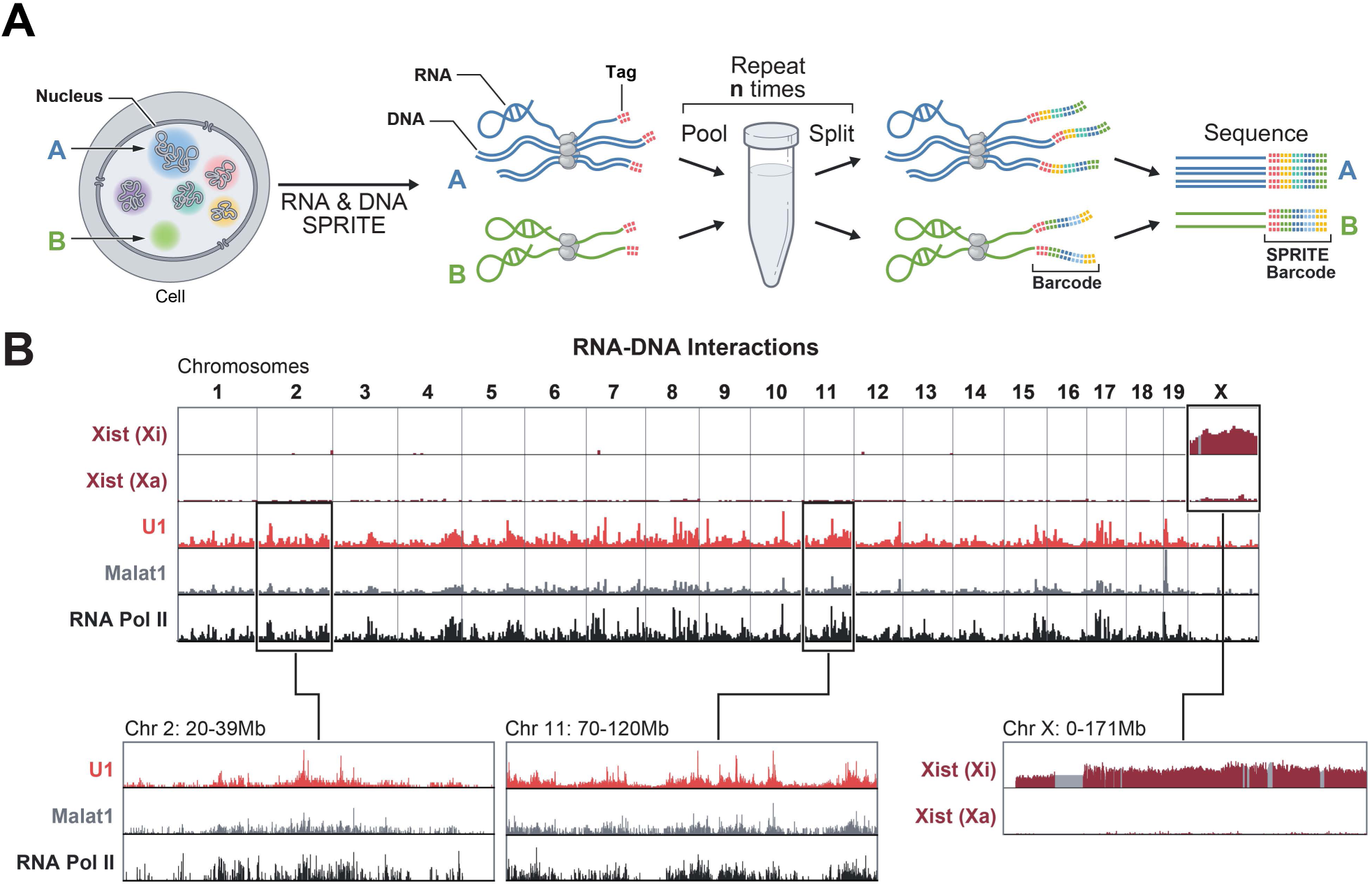
RD-SPRITE generates maps of higher-order RNA and DNA contacts throughout the cell. **(A)** Schematic of the RD-SPRITE protocol. Crosslinked cells are fragmented into smaller crosslinked complexes (e.g. A, B). RNA and DNA are each tagged with a DNA-specific or RNA-specific adaptor sequence (pink). The sample is processed through multiple rounds of split-and-pool barcoding (*n* times), where tag sequences are concatemerized during each round. A series of tags is referred to as a SPRITE barcode. RNA and DNA are sequenced, and barcodes are matched to generate SPRITE clusters to identify all interacting molecules. **(B)** RNA-DNA interactions of various non-coding RNAs. Xist (burgundy) unweighted contacts across the genome in female ES cells where Xist is induced exclusively on the 129 allele (Xi), but not the Castaneous allele (Xa). U1 spliceosomal RNA (red) and Malat1 lncRNA (grey) weighted contacts across the genome occur at highly transcribed RNA Pol II (ENCODE) genomic regions (black). Insets show zoom-ins of Xist (right) and U1/Malat1 along with genomic localization of RNA Pol II from ENCODE (middle and left). Masked regions on chromosome X plotted in gray.

We performed RD-SPRITE in an F1 hybrid female mouse ES cell line that was engineered to induce Xist from a single allele (see **Methods**). We sequenced these libraries on a NovaSeq S4 run to generate ∼8 billion reads corresponding to ∼720 million SPRITE clusters (**Supplemental Figure 1C**). We confirmed that we accurately identify RNA- and DNA-specific reads (**Supplemental Figure 1A-B**) and that the data measure *bona fide* RNA interactions – including well-described RNA-DNA and RNA-RNA contacts not only in the nucleus, but throughout the cell.

First, we explored RNA-DNA contacts captured in our data and compared their interactions to those of several ncRNAs that have been previously mapped to chromatin that reflect a range of known *cis* and *trans* localization patterns. Specifically, we observed strong enrichment of: **(i)** Xist over the inactive X (Xi), but not the active X chromosome (Xa)^46,56^ (**Figure 1B, Supplemental Figure 1D**); **(ii)** Malat1 and U1 over actively transcribed RNA Polymerase II genes^57,58^ (**Figure 1B**); and **(iii)** telomerase RNA component (Terc) over telomere-proximal regions of all chromosomes (**Supplemental Figure 1E**)^59,60^.

Second, we explored known RNA-RNA contacts that occur in different locations in the cell. For example, we observed a large number of contacts between translation-associated RNAs in the cytoplasm, including all RNA components of the ribosome (5S, 5.8S, 18S, 28S) and ∼8000 individual mRNAs (exons), but not with pre-mRNAs (introns). Conversely, we observed many contacts between the small nuclear RNA (snRNA) components of the spliceosome (e.g. U1, U2, U4, U5, U6) in the nucleus and individual pre-mRNAs (**Figure 2A**).

**Figure 2:**
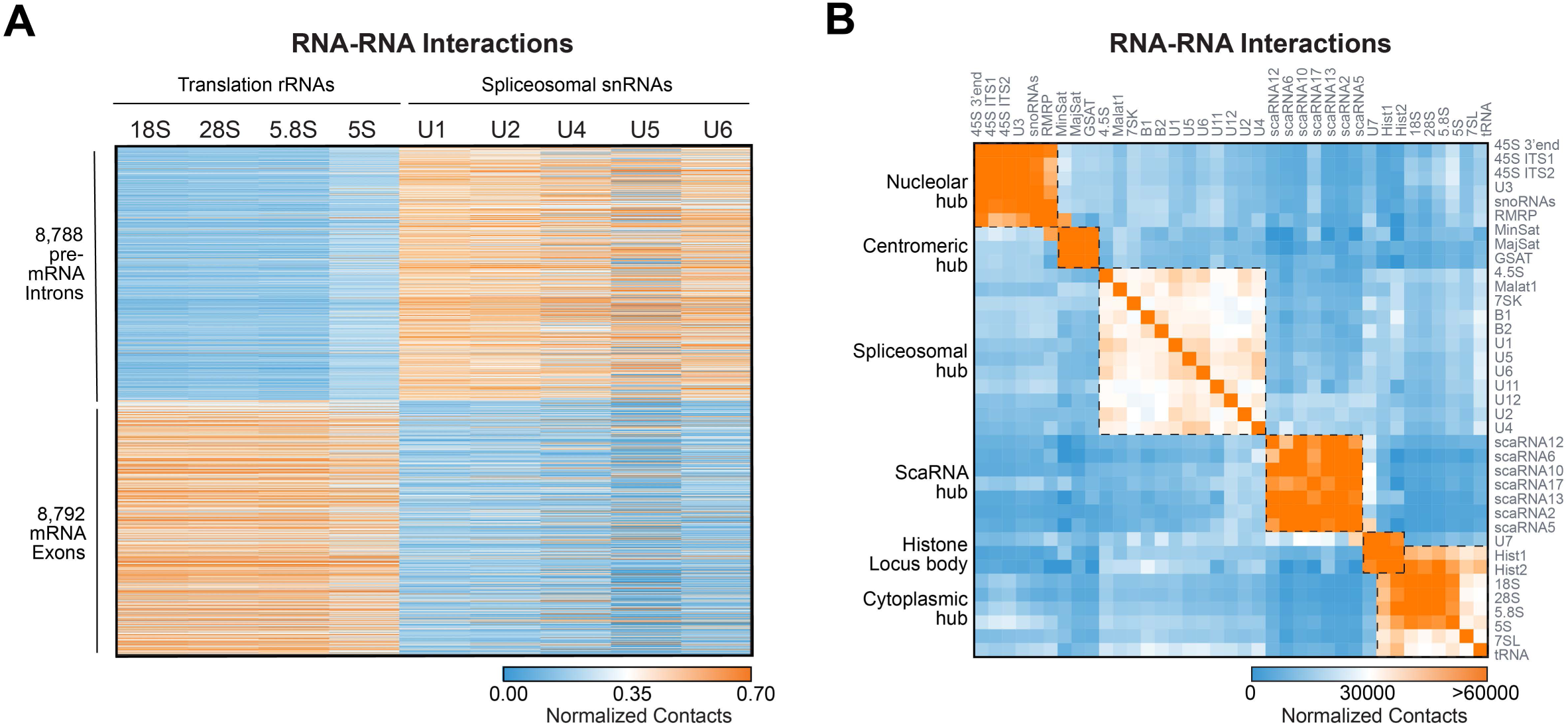
Non-coding RNAs form hubs containing RNAs of shared functional roles in different cellular locations. **(A)** A heatmap showing the number of unweighted RNA-RNA contacts between different classes of RNAs. Columns: translation-associated RNAs (18S, 28S, 5.8S, and 5S) and splicing-associated RNAs (U1, U2, U4, U5, U6). Rows: Introns and exons of individual mRNAs. Orange represents high contact frequency and blue represents low contact frequency. **(B)** A heatmap showing RNA-RNA unweighted contact frequencies for several classes of RNAs. Orange represents high contact frequency and blue represents low contact frequency. Groups of RNAs that have high contact frequencies with each other, but not other RNAs, are referred to as RNA hubs.

Together, these results demonstrate that RD-SPRITE accurately measures known RNA-DNA and RNA-RNA localization patterns in the nucleus and cytoplasm. While we focus primarily on RNA localization within the nucleus, we note that RD-SPRITE can also be utilized to study RNA compartments beyond the nucleus^61–63^.

### Non-coding RNAs localize in higher-order spatial compartments in the nucleus

Because RD-SPRITE generates comprehensive structure maps of RNA and DNA in the nucleus, we explored which specific RNAs localize within higher-order compartments. To do this, we mapped all RNA-RNA and RNA-DNA interactions genome-wide. Specifically, we identified several sets of RNA and genomic DNA regions that display high contact frequencies within their corresponding set, but low contact frequencies with molecules contained within distinct sets. We refer to the RNA and DNA molecules within an interacting set as an RNA-DNA hub (**Figure 2B, Supplemental Figure 2A-B**). Using a combination of RNA FISH (to visualize RNAs) and immunofluorescence (to visualize different cellular compartments) we confirmed that RNAs within a hub co-localize (**Supplemental Figure 2C**), while RNAs in distinct hubs localize to different regions of the cell (**Supplemental Figure 2D**). Using multi-way SPRITE clusters, we found that these RNA-DNA hubs form higher-order structures in the nucleus that contain multiple RNAs and genomic DNA regions that are organized in 3D space (**Figure 3B**) around shared regulatory functions, which we describe below.

**Figure 3:**
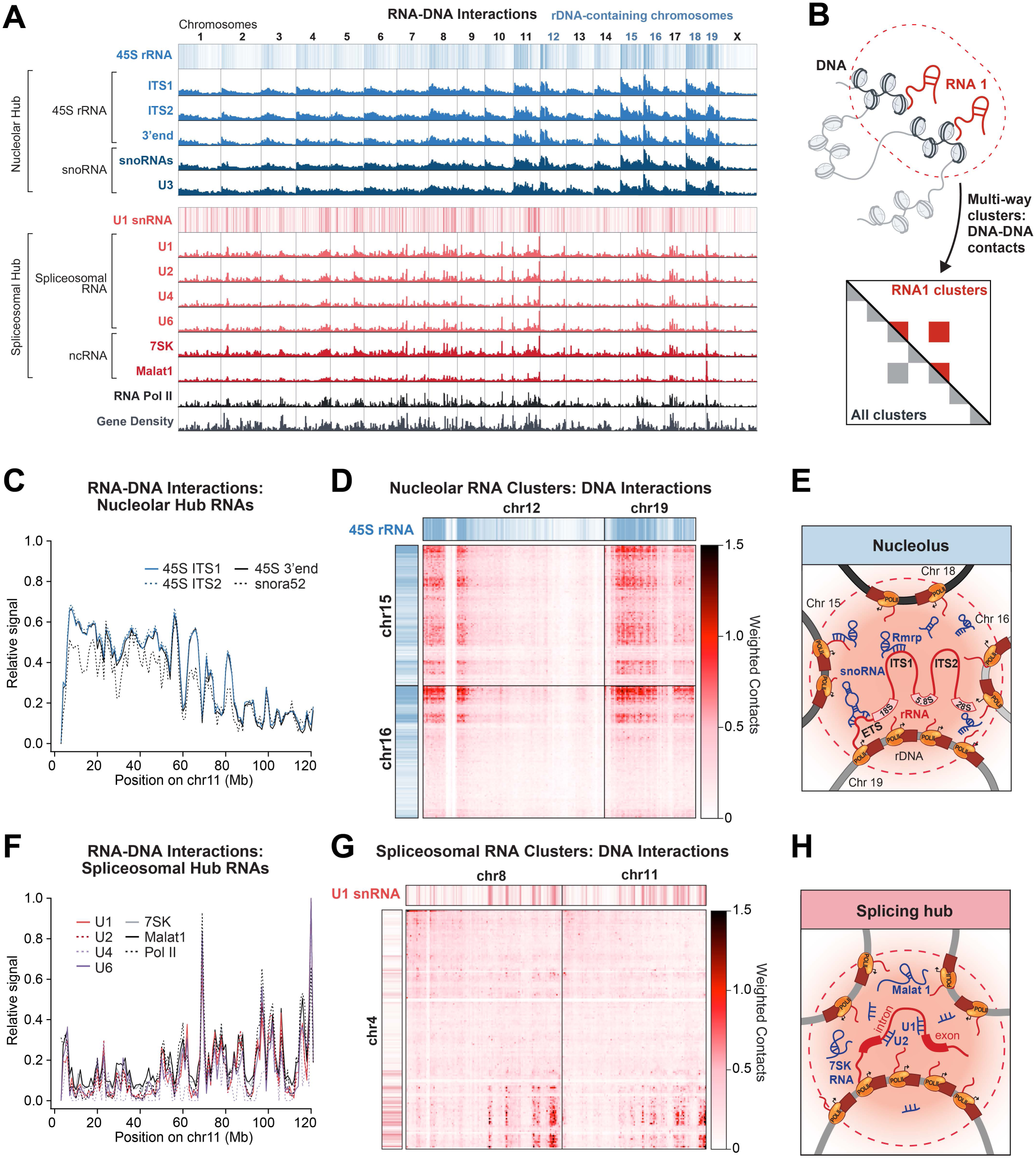
Nucleolar and spliceosomal RNAs form genome-wide interaction hubs. **(A)** Genome-wide weighted RNA-DNA contacts (1Mb resolution) for several RNAs within the nucleolar (blue) and spliceosomal (red) hubs. RNA Pol II occupancy from ENCODE is shown along with gene density across the genome. Chromosomes that contain genes for ribosomal RNA, which are located at the centromere proximal regions of each chromosome, are demarcated in blue (chr. 12, 15, 16, 18, and 19). Blue and red horizontal heatmaps represent RNA-DNA interactions of the 45S (3’end) pre-rRNA and U1 snRNA (1Mb resolution). (**B**) SPRITE can measure the 3D organization of DNA occurring within RNA hubs. Multi-way RNA-DNA clusters can be used to measure DNA-DNA contacts (illustrative heatmap) occurring specifically at DNA loci interacting with a given RNA (RNA1 clusters, red contacts) or across all SPRITE clusters (All clusters, gray contacts), as shown in the upper half and lower half of the diagonal, respectively. **(C)** Overlay of RNA-DNA contact frequencies on chromosome 11 is shown for various RNAs within the nucleolar hub. **(D)** Weighted DNA-DNA contacts within SPRITE clusters containing nucleolar hub RNAs (e.g 45S pre-rRNAs, snoRNAs, Rmrp). Long range, higher-order inter-chromosomal nucleolar interactions are shown between chromosomes 12 and 19 and chromosomes 15 and 16 for nucleolar hub RNA-containing clusters. **(E)** Schematic of our observations showing nascent pre-rRNAs in spatial proximity to their loci and snoRNAs and Rmrp binding to these RNAs that are spatially concentrated within this compartment. **(F)** Overlay of RNA-DNA contact frequencies on chromosome 11 is shown for the spliceosomal hub examples above. **(G)** Weighted DNA-DNA contacts within SPRITE clusters containing spliceosomal hub RNAs (e.g. U1, U2, Malat1, 7SK). Long range, higher-order inter-chromosomal spliceosomal hub interactions are shown between regions on chromosome 4 and chromosomes 8 and 11 (examples that have high Pol II occupancy) for all spliceosomal hub RNA-containing clusters. **(H)** Schematic of our observation showing nascent pre-mRNAs in spatial proximity to their transcriptional loci and snRNAs (e.g. U1 and U2 shown) that bind to pre-mRNAs that, along with Malat1 and 7SK, are spatially concentrated within this compartment around transcribed genomic DNA.

### Non-coding RNAs form processing hubs around genomic DNA encoding their nascent targets

We first explored the RNA-DNA hubs corresponding to several nuclear compartments associated with RNA processing. Biochemical approaches have revealed that RNA processing generally involves direct hybridization between various diffusible *trans*-acting ncRNAs (e.g. splicing snRNAs) and their nascent target RNA substrates (e.g. pre-mRNAs). However, these approaches study a few molecules at a time and require a priori knowledge of which molecular components comprise specific nuclear bodies. Using RD-SPRITE to measure the higher-order organization of RNA and DNA molecules within each processing hub, we examined: (i) the RNA components in these hubs (RNA-RNA interactions), (ii) the location of each RNA relative to the DNA loci from which their nascent target substrate RNAs are transcribed (RNA-DNA interactions), and (iii) whether RNA processing occurs at individual locations or whether multiple DNA loci come together in 3D space (DNA-DNA interactions).

#### (i) ncRNAs involved in ribosomal RNA processing organize within a 3D compartment containing transcribed ribosomal RNA genes

We identified a hub that includes the 45S pre-ribosomal RNA (pre-rRNA), RNase MRP, and dozens of snoRNAs that are involved in ribosomal RNA biogenesis (**Figure 2B, Supplemental Figure 2A, 3A**). rRNA is transcribed as a single 45S precursor RNA and is cleaved by RNAse MRP and modified by various snoRNAs to generate the mature 18S, 5.8S, and 28S rRNAs^64–66^. We found that all of these RNAs diffuse through the nucleus and localize at genomic locations that are proximal to ribosomal DNA repeats that encode the 45S pre-rRNA and other genomic regions that we previously showed to organize around the nucleolus^54^ (**Figure 3A, 3C**, see **Methods**). We explored the DNA-DNA interactions that occur within SPRITE clusters containing multiple nucleolar hub RNAs (45S pre-rRNA and snoRNAs, ≥4-way contacts) (**Figure 3B**), and observed that these RNAs and the multiple genomic DNA regions encoding 45S pre-RNAs are organized together in 3D space (**Figure 3D, Supplemental Figure 3B**, see **Methods**). Our results demonstrate that the nascent 45S pre-rRNA is enriched near the DNA loci from which it is transcribed. In this way, 45S pre-rRNA (which is known to directly interact with snoRNAs and RNase MRP^22,64^) may act to concentrate these diffusible *trans*-acting regulatory ncRNAs that are responsible for ribosome biogenesis into the nucleolar compartment (**Figure 3E**). Consistent with this, inhibition of 45S pre-rRNA transcription has been shown to disrupt nucleolar organization^9,42,43^.

#### (ii) ncRNAs involved in mRNA splicing are spatially concentrated around transcribed Pol II genes

We identified a hub that contains nascent pre-mRNAs along with all of the major (e.g. U1, U2, U4, U5, U6) and minor (U11, U12) spliceosomal ncRNAs and other ncRNAs associated with transcriptional regulation and mRNA splicing (e.g. 7SK and Malat1) (**Figure 2A-B**). Nascent pre-mRNAs are known to be directly bound and cleaved by spliceosomal RNAs to generate mature mRNA transcripts^24,67^. Although splicing can occur co-transcriptionally^24,68,69^, it has been unclear how spliceosomal RNAs are organized in the nucleus relative to target pre-mRNAs and genomic DNA^69–74^. We found that the spliceosomal hub RNAs localize to genomic DNA regions containing actively transcribed Pol II genes (Pearson r = 0.84-0.90, **Figure 3A, 3F, Supplemental Figure 3C**). We explored DNA-DNA contacts within SPRITE clusters containing individual and multiple spliceosomal hub RNAs (≥2 distinct RNAs, ≥4-way RNA-DNA contacts) and observed that these RNAs and genomic DNA regions form preferential intra- and inter-chromosomal contacts that are organized together in 3D space (**Figure 3G, Supplemental Figure 3D**). These results demonstrate that spliceosomal RNAs are spatially organized around clusters of actively transcribed Pol II genes and their associated nascent pre-mRNAs (**Figure 3H**). Because nascent pre-mRNAs are enriched in spatial proximity to their transcriptional locus and are known to directly hybridize to splicing RNAs^75,76^, nascent pre-mRNAs may act to recruit these diffusible *trans*-acting regulatory ncRNAs into high spatial concentrations near their co-transcriptional targets.

#### (iii) ncRNAs involved in snRNA biogenesis are spatially organized around snRNA gene clusters

We identified a hub containing several annotated small Cajal body-associated RNAs (scaRNAs), two previously unannotated scaRNAs, and several small nuclear RNAs (snRNAs) (**Figure 2B, Supplemental Figure 4D**, see **Methods**). snRNAs are Pol II transcripts produced from multiple locations throughout the genome that undergo 2’O-methylation and pseudouridylation before functionally acting as components of the spliceosome at thousands of nascent pre-mRNA targets^77–79^; scaRNAs directly hybridize to snRNAs to guide these modifications^80–82^. We found that scaRNAs are highly enriched at discrete genomic regions containing multiple snRNA genes in close linear space (**Figure 4A**). Despite being separated by large genomic distances, these DNA regions form long-range contacts in SPRITE clusters containing scaRNAs (≥3-way RNA-DNA contacts) (**Figure 4B, Supplemental Figure 4E**). In fact, we observe that these scaRNAs, snRNAs, and the distal DNA loci from which the snRNAs are transcribed simultaneously interact within higher-order SPRITE clusters, demonstrating that all of these components interact within a 3D hub in the nucleus (**Supplemental Figure 4G**). Because snRNAs are enriched in spatial proximity to their transcriptional loci and are known to directly hybridize to diffusible scaRNAs^81,82^, nascent snRNAs may act to recruit and concentrate scaRNAs within this compartment to enable snRNA biogenesis and modification^83^ (**Figure 4E**). We note that this snRNA biogenesis hub may be similar to Cajal bodies, which have been noted to contain snRNA genes and scaRNAs^76,84–87^ (see **Supplementary Note 2**).

**Figure 4:**
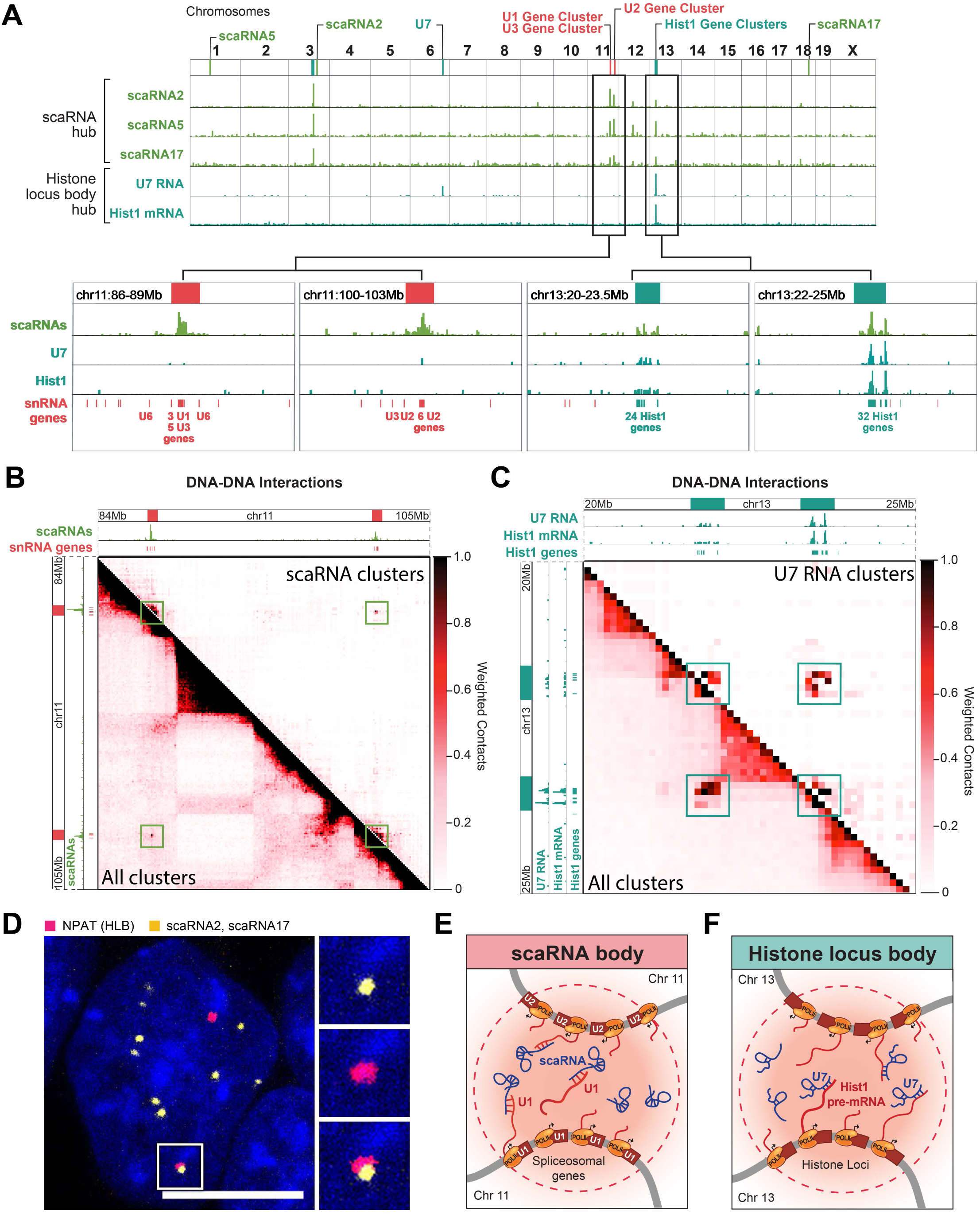
Non-coding RNAs involved in snRNA and histone mRNA biogenesis are organized around snRNA and histone gene clusters. **(A)** Weighted RNA-DNA contacts for scaRNA2, scaRNA5 (Gm25395), and scaRNA17 (green) and U7 RNA and histone pre-mRNAs (teal) are plotted across the genome. Insets (bottom) show zoom-ins on specific regions. Lines (top) show genomic locations of each RNA plotted and gene cluster of interest. **(B)** Weighted DNA-DNA contacts within SPRITE clusters containing scaRNAs (upper diagonal) along with all weighted DNA-DNA contacts within all SPRITE clusters (lower diagonal) are shown across a region of chromosome 11. The locations of scaRNA occupancy is shown along the top and side axes and demarcated by a solid red box. **(C)** Weighted DNA-DNA contacts within SPRITE clusters containing the U7 RNA (upper diagonal) along with all weighted DNA-DNA contacts within all SPRITE clusters (lower diagonal) are shown across a region of chromosome 13. U7 and histone occupancy is shown along the top and side axis and demarcated with a teal box. **(D)** RNA FISH of scaRNAs (pooled scaRNA2 and scaRNA17 probes) along with IF of a known histone locus body (HLB) marker (NPAT) show proximity of the two compartments. Scalebar is 10μm. **(E)** Schematic showing scaRNAs spatially concentrated near the transcribed genomic loci of target snRNA gene clusters. **(F)** Schematic showing U7 RNA spatially concentrated near the transcribed genomic loci of target histone mRNA gene loci.

#### (iv) The histone processing U7 snRNA is spatially enriched around histone gene loci

We identified a hub containing the U7 snRNA and various histone mRNAs (**Figure 2B**). Unlike most pre-mRNAs, histone pre-mRNAs are not polyadenylated; instead their 3’ends are bound and cleaved by the U7 snRNP complex to produce mature histone mRNAs^31,88^. This process is thought to occur within nuclear structures called Histone Locus Bodies (HLBs)^32,80^. We observed that the U7 snRNA localizes at genomic DNA regions containing histone mRNA genes, specifically, at two histone gene clusters on chromosome 13 (**Figure 4A**). To determine whether the U7 snRNA, histone gene loci, and nascent histone pre-mRNAs form a 3D spatial compartment, we generated DNA-DNA interaction maps from U7 snRNA-containing clusters (≥3-way RNA-DNA contacts) and observed long-range DNA contacts between the two histone gene clusters on chromosome 13 (**Figure 4C, Supplemental Figure 4F**). Because histone pre-mRNAs are present at high concentrations near their transcriptional loci and directly bind to U7, they may act to recruit these *trans*-associating ncRNAs into the HLB compartment (**Figure 4F**). Consistent with this model, previous studies have shown that histone pre-mRNAs are sufficient to seed the formation of the HLB and that the U7 binding site on the histone pre-mRNA is required for HLB formation^80,89,90^. Additionally, we observed that scaRNAs also localize to these histone gene clusters, form higher-order DNA interactions, and are adjacent to the HLB in the nucleus (**Figure 4A**,**D**, **Supplemental Figure 4C**,**H**). This is consistent with previous observations that HLBs and Cajal bodies are often found adjacent to each other in the nucleus^80,85^ and tethering histone pre-mRNAs to chromatin can also lead to recruitment of components of Cajal bodies^89^.

These results demonstrate that RD-SPRITE comprehensively and simultaneously maps RNA and DNA molecules comprising several well-defined RNA processing hubs. Specifically, in all of these cases, we observe that: (i) nascent substrate RNAs interact with diffusible *trans*-associating ncRNAs (RNA-RNA), (ii) these nascent RNAs are localized near their DNA loci (RNA-DNA), and (iii) multiple DNA loci containing these nascent transcripts come together in 3D space (DNA-DNA). Using SPRITE, we previously detected higher order DNA organization around two landmark nuclear bodies – nucleoli and nuclear speckles^54^; our results now show that higher-order organization of multiple DNA loci (DNA-DNA) around shared targets is a general principle of RNA processing that includes ribosomal RNA, mRNA, snRNA, and histone mRNA biogenesis.

### Satellite-derived ncRNAs organize HP1 localization at inter-chromosomal hubs

In addition to RNA processing, we also identified a hub containing ncRNAs transcribed from minor and major satellite DNA regions within centromeric and pericentromeric regions, respectively (**Figure 2B**). We found that these ncRNAs localize primarily over centromere-proximal regions (**Figure 5A-B, Supplemental Figure 5B**) and organize into higher-order structures containing these ncRNAs and multiple centromere-proximal regions from different chromosomes that interact simultaneously (**Figure 5C, Supplemental Figure 5A**). This suggests that these RNAs demarcate a nuclear body where centromeric regions of chromosomes interact with each other. To confirm this, we performed DNA FISH on the major and minor satellite DNA and observed higher-order structures where multiple centromeres from distinct chromosomes interact simultaneously^91,92^ (**Figure 5D**).

**Figure 5:**
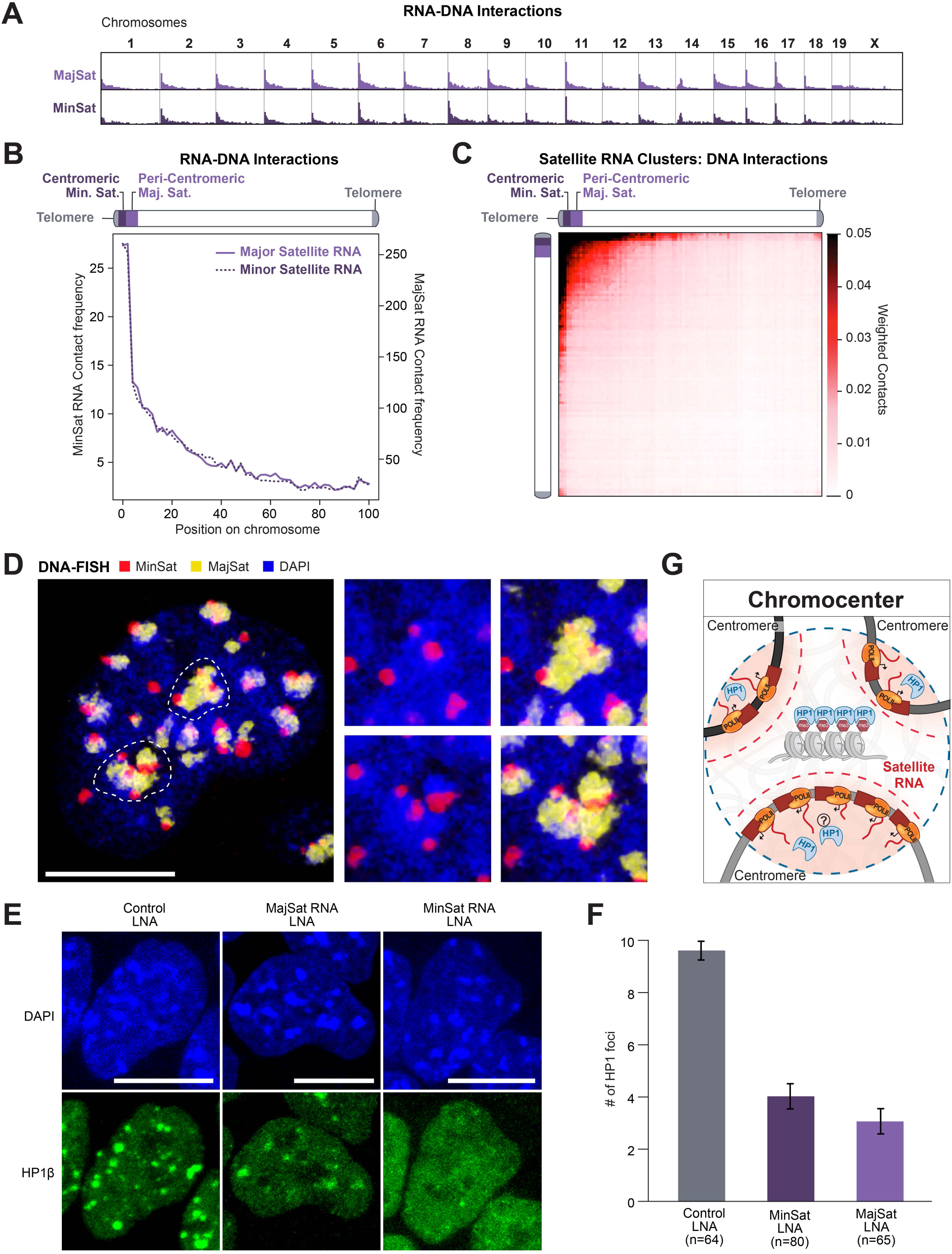
Satellite-derived ncRNAs organize HP1 localization at inter-chromosomal hubs. **(A)** Unweighted RNA-DNA contact frequencies of major (dark purple) and minor (light purple) satellite-derived ncRNAs on DNA. **(B)** Aggregate unweighted contact frequencies of major and minor satellite-derived RNAs on DNA across all chromosomes. **(C)** Weighted inter-chromosomal DNA-DNA contacts within SPRITE clusters containing satellite-derived RNAs. **(D)** DNA FISH on major (yellow) and minor (red) satellite DNA shows multiple centromeres organize around distinct chromocenter structures within a nucleus (DAPI). Dashed lines demarcate two chromocenter structures shown on the right. Specifically, we observe multiple individual centromeres (visualized by minor satellite DNA) and the larger peri-centromeric regions (visualized by major satellite DNA) organized at the focal DAPI-dense chromocenter structures. Scalebar is 10μm. **(E)** LNA-mediated knockdown of major and minor satellite-derived RNA (referred to as MajSat and MinSat RNA, respectively). (Left) Control LNA knockdown and HP1β immunofluorescence. (Middle) Disruption of HP1β foci with LNA knockdown of MajSat RNA. (Right) Disruption of HP1β foci with LNA knockdown of MinSat RNA. Scalebar is 10μm. **(F)** Quantification of the mean number of HP1 foci per cell in (E). HP1β foci above a given intensity threshold were quantified (see Methods). Violin plot provided in Supplemental Figure 5E. Control: n=64 cells, MinSat: n=80 cells, MajSat: n=65 cells. Error bars represent standard error. **(G)** Schematic showing satellite RNAs (red gradient) spatially concentrated near the pericentromeric and centromeric DNA around a heterochromatic chromocenter structure (center), which is highly enriched with HP1 protein.

Higher-order organization of centromeric and pericentromeric DNA, often referred to as chromocenters, represent one of the most well-defined regions of heterochromatin assembly and are enriched for various heterochromatin enzymes and chromatin modifications, including the HP1 protein and H3K9me3 modifications^92^. Previous studies have shown that global disruption of RNA by RNase A leads to disruption of HP1 localization at chromocenters^91^. However, RNAse A is not specific and can impact several structures in the nucleus, including nucleoli^93^. Because major and minor satellite-derived ncRNAs localize exclusively within centromere-proximal structures, we hypothesized that these ncRNAs might be important for HP1 localization. To test this, we used a locked nucleic acid (LNA) antisense oligonucleotide (ASO) to degrade either the major or minor satellite RNAs (see **Methods**). We found that disruption of either the major or minor satellite RNA leads to depletion of HP1 proteins over these centromere-proximal structures and altered chromocenter organization (**Figure 5E-F, Supplemental Figure 5C-F**, see **Supplemental Note 4**). Our results demonstrate that major and minor satellite RNAs are enriched within spatial proximity of their transcriptional loci and are required to recruit HP1 into centromere-proximal nuclear compartments to maintain higher-order centromeric heterochromatin (**Figure 5G**). Consistent with this, previous studies have shown that disruption of the major satellite-derived RNA prior to the formation of chromocenters during preimplantation development leads to loss of chromocenter formation, lack of heterochromatin formation, and embryonic arrest^94–96^.

### Hundreds of non-coding RNAs localize in spatial proximity to their transcriptional loci

Thousands of nuclear-enriched ncRNAs are expressed in mammalian cells, but only a handful have been mapped on chromatin. We mapped ∼650 lncRNAs in ES cells and observed a striking difference in chromatin localization between these lncRNAs and mature mRNAs (**Figure 6A, Supplemental Figure 6A-B**, see **Supplemental Note 3**). Specifically, we found that the vast majority (93%) of these lncRNAs are strongly enriched within 3D proximity of their transcriptional loci (**Figure 6B-D, Supplemental Figure 6C**, see **Methods**). This is in contrast to mature mRNAs, which are depleted near their transcriptional loci and at all other genomic locations (**Supplemental Figure 6D-E**). We observed a similar lack of chromatin enrichment for a subset of lncRNAs (enrichment score <0), including Norad which is known to localize and function in the cytoplasm^97^ (**Figure 6A-B**). Additionally, not all lncRNAs with high chromatin enrichment are restricted to the 3D compartment around their locus. For example, the Malat1 lncRNA is strongly enriched on chromatin but localizes broadly across all chromosomes (**Figure 6A-B, Supplemental Figure 6C**). These data demonstrate that the vast majority of lncRNAs in ES cells localize exclusively in spatial proximity to their transcriptional loci and do not diffuse to other locations in the nucleus or cytoplasm (**Figure 6D-E**).

**Figure 6:**
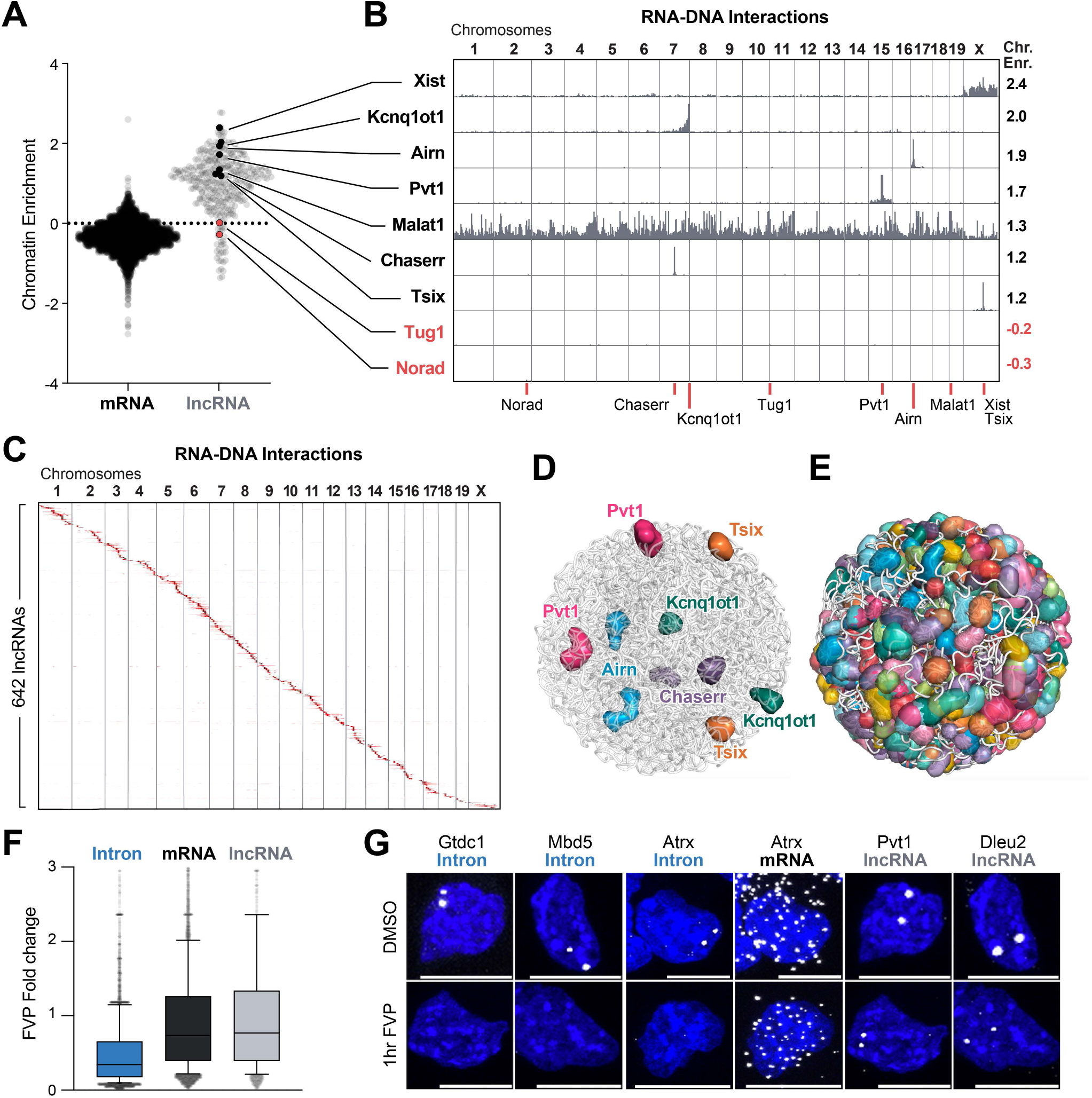
Most lncRNAs localize at genomic targets in 3D proximity to their transcriptional loci. **(A)** Chromatin enrichment score for mRNAs (black) and lncRNAs (gray). Values greater than 0 represent RNAs enriched on chromatin versus values less than 0 represent RNAs depleted on chromatin. Chromatin enrichment scores for all classes of RNAs are provided in Supplemental Figure 6A-B. **(B)** Unweighted RNA-DNA localization maps across the genome for selected chromatin enriched (black) and chromatin depleted (red) lncRNA examples. Chromatin enrichment scores (right) for each lncRNA are listed. Red lines (bottom) show genomic locations of each RNA plotted. **(C)** Unweighted genome-wide RNA-DNA localization map of 642 lncRNAs (rows) ordered by the genomic position of their transcriptional loci. **(D)** A 3D space filling nuclear structure model of the selected lncRNAs shown in (B). **(E)** A 3D space filling nuclear structure model based of 543 lncRNAs that display at least 50-fold enrichment in the nucleus. Each sphere corresponds to a 1 Mb region or larger where each lncRNA is enriched. **(F)** Analysis of global run on sequencing (GRO-seq) data from Jonkers *et. al*. (2014)^99^ comparing the fold change in RNA levels (TPMs) between untreated and 50 minutes of flavopiridol (FVP) in mouse ES cells. Changes in introns (blue), mRNAs (black), and lncRNAs (gray) are shown. Box and whiskers plot represents median, box extends from 25th to 75th percentiles, and whiskers are drawn from the 10th to 90th percentiles. **(G)** RNA FISH for selected introns (Gtdc1, Mbd5, Atrx), mRNA exons (Atrx), and lncRNAs (Pvt1, Dleu2) treated for 1 hour with DMSO or FVP are shown. Scalebar is 10μm.

This exclusive localization pattern could reflect the formation of a stable RNA-enriched nuclear compartment or simply represent an unstable RNA product that is transiently associated with its transcriptional locus prior to being rapidly degraded (**Supplemental Figure 6A**). To exclude the possibility that these represent unstable RNA products, we explored the expression of these lncRNAs after treating cells with flavopiridol (FVP), a drug that runs off elongating Pol II and prevents re-initiation of transcription^98^. We explored a previously published global RNA sequencing experiment performed after 50 minutes of treatment with FVP in mES cells^99^. Consistent with previous reports^100^, we found that virtually all lncRNAs were dramatically more stable than nascent pre-mRNAs and comparable to the stability of mature mRNAs (**Figure 6F**). To confirm this, we performed RNA FISH for 4 lncRNAs, 6 nascent pre-mRNAs (introns), and 1 mature mRNA (exons) in untreated cells and upon FVP treatment (see **Methods**). We found that all of the lncRNAs form stable nuclear foci that are retained upon transcriptional inhibition (**Figure 6G, Supplemental Figure 6F**). In contrast, all nascent pre-mRNA foci are lost upon transcriptional inhibition, even though we observe no impact on their mature mRNA products (**Figure 6G**).

Together, these results demonstrate that many lncRNAs form high concentration territories within defined nuclear compartments. While their complete molecular composition and functional relevance remain unclear, our results demonstrate that these RNA-compartments are widespread and can demarcate local spatial territories throughout the nucleus (**Figure 6E**).

### Non-coding RNAs guide regulatory proteins to nuclear compartments to regulate gene expression

Because hundreds of lncRNAs are enriched in spatial compartments throughout the nucleus, we explored how RNA localization might impact protein localization within these compartments. Recently, we and others showed that SHARP (also called Spen) directly binds Xist^101–105^ and recruits the HDAC3 histone deacetylase complex to the X chromosome to silence transcription^103,106,107^ (**Supplemental Figure 7A**). To explore the nuclear localization of SHARP more globally, we performed super-resolution microscopy and found two types of SHARP localization: low-level diffuse localization throughout the nucleus and compartmentalized localization within dozens of well-defined, high-intensity, foci throughout the nucleus (∼50-100 foci/nucleus) (**Figure 7B**, see **Supplemental Video 1**). To determine whether the compartmentalized SHARP foci are dependent on RNA, we deleted the RNA binding domains (ΔRRM) from the protein (**Figure 7A**) and found that it led to loss of all compartmentalized SHARP foci without affecting diffusive localization of the protein throughout the nucleus (**Figure 7B**, see **Supplemental Video 2**). These results demonstrate that RNA is required for SHARP localization to dozens of compartments throughout the nucleus.

**Figure 7:**
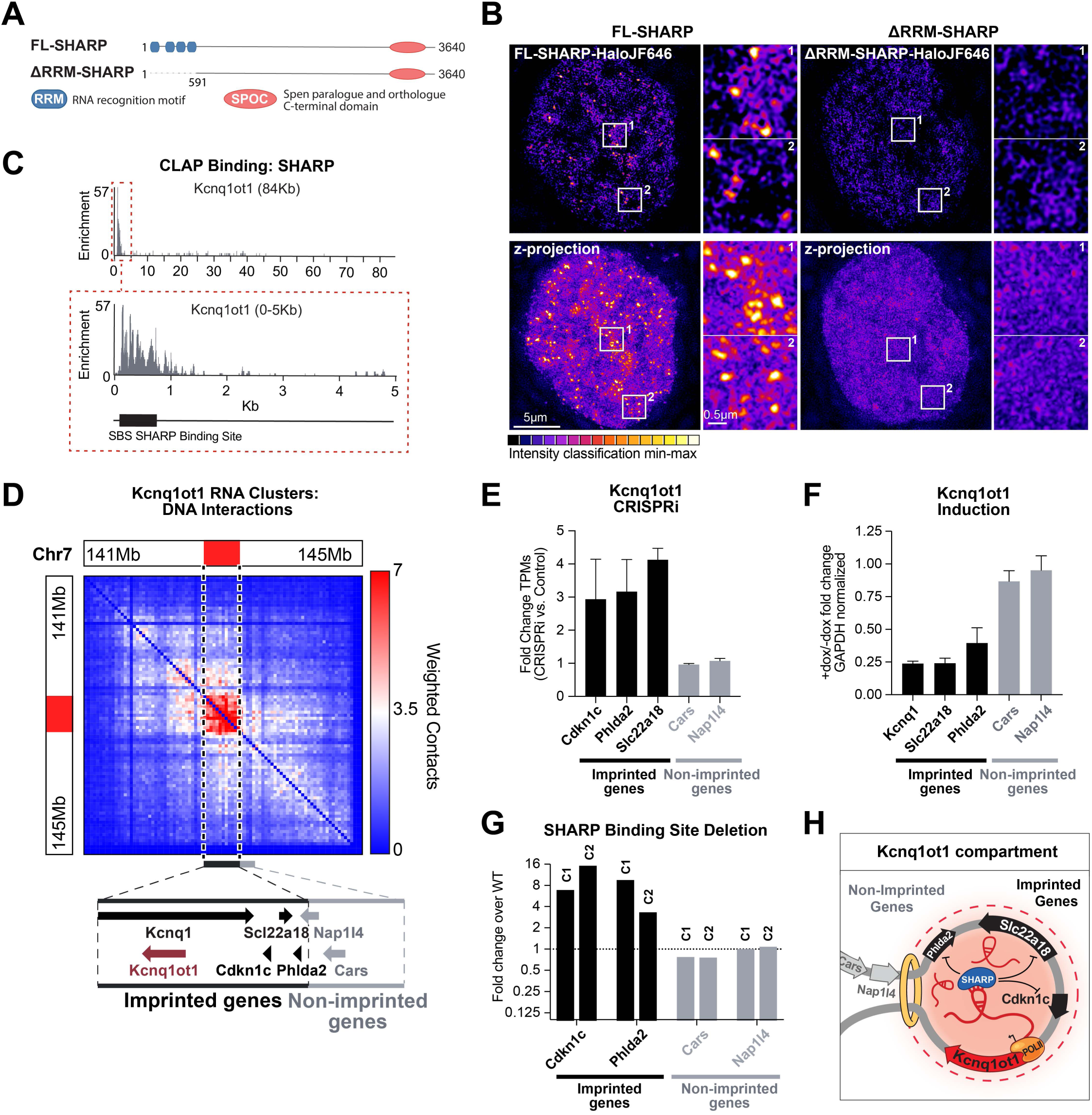
Kcnq1ot1 seeds an RNA-mediated compartment over its imprinted target genes and binds the SHARP protein to repress gene expression. **(A)** Diagram of the functional domains in full length (FL) SHARP (also referred to as Spen) protein. SHARP contains four RNA recognition motif (RRM, blue) domains and one Spen paralogue and orthologue C-terminal (SPOC, orange) domain. A version of SHARP lacking its RNA binding motifs (ΔRRM) was generated by deletion of the first 591 amino acids of SHARP. **(B)** Top 3D-SIM 125 nm optical sections of FL-SHARP (left) and ΔRRM-SHARP (right) and z-projections (bottom) Halo-tagged FL- and ΔRRM-SHARPJF646. FL-SHARP localizes in foci throughout the nucleus (zoom in panels 1-2), while ΔRRM-SHARP leads to diffusive localization. Bar: 5μm, insets: 0.5μm (magnifications). Intensities are depicted in 16-color grading from black (minimum) to white (maximum). **(C)** SHARP (also referred to as Spen) protein binding on the entire 84,000 nucleotide Kcnq1ot1 lncRNA (top), and a zoom in on the first 5,000 nucleotides of the lncRNA (bottom). We define a region called the SHARP binding site (SBS) shown as a black box. **(D)** Weighted DNA-DNA contacts within SPRITE clusters containing the Kcnq1ot1 RNA. Dashed line indicates the location of the Kcnq1ot1-enriched spatial compartment. (Zoom out) Genomic locations in this domain of the Kcnq1ot1 gene (burgundy) and imprinted target genes Kcnq1, Slc22a18, Cdkn1c, and Phlda2 (black) and non-imprinted neighboring genes Nap1l4 and Cars (gray). **(E)** Changes in mean gene expression upon CRISPR inhibition (CRISPRi) of the Kcnq1ot1 lncRNA. Genes contained within the Kcnq1ot1-associated domain (e.g. Cdkn1c, Phlda2, Slc22a18) are shown in black and genes outside the domain (e.g. Cars, Nap1l4) are shown in gray. Error bars represent standard deviation. **(F)** Changes in mean gene expression upon doxycycline mediated induction (+Dox) of Kcnq1ot1 relative to cells with no doxycycline (-Dox). Genes contained within the Kcnq1ot1-associated domain (e.g. Cdkn1c, Phlda2, Slc22a18) are shown in black and genes outside the domain (e.g. Cars, Nap1l4) are shown in gray. Error bars represent standard deviation. **(G)** Homozygous deletion of the SHARP binding site (**Δ**SBS) in two different clones results in up-regulation of the genes within the Kcnq1ot1-domain (e.g. Cdkn1c, Phlda2), but does not impact the neighboring genes outside this domain (e.g. Cars, Nap1l4). **(H)** Schematic of our results for the Kcnq1ot1-mediated compartment. Kcnq1ot1 lncRNA seeds the formation of an RNA-mediated compartment in spatial proximity to its transcriptional locus, and then binds and recruits the SHARP protein into this compartment to silence its imprinted target genes.

To explore how these ncRNA-mediated nuclear compartments might act to regulate gene expression, we purified SHARP and mapped its interactions with specific RNAs. We identified strong binding to several RNAs, including a ∼600 nucleotide region at the 5’ end of Kcnq1ot1, a lncRNA associated with the pediatric Beckwith-Wiedemann overgrowth syndrome^108^ (see **Methods, Figure 7C**). We found that Kcnq1ot1 localizes within the topologically associating domain (TAD) that contains all of the known paternally-imprinted genes (Cdkn1c, Slc22a18, Phlda2)^108,109^, but excludes other genes that are close in linear space in the genome (e.g. Cars, Nap1l4, **Figure 7D, Supplemental Figure 7G**). We confirmed that downregulation of Kcnq1ot1 using CRISPRi leads to upregulation of these target genes and, conversely, bi-allelic induction of Kcnq1ot1 expression leads to silencing of these imprinted target genes. In both cases, there was no impact on the genes outside of this Kcnq1ot1-associated domain (**Figure 7E-F, Supplemental Figure 7C**).

To explore whether SHARP binding to the Kcnq1ot1 RNA is essential for transcriptional silencing in this compartment, we deleted the SHARP binding site on Kcnq1ot1 (ΔSBS) and observed upregulation of its known target genes in two independent clones (**Figure 7G**). Because SHARP is known to recruit HDAC3^103,106,107^, we tested whether Kcnq1ot1-mediated silencing is dependent on histone deacetylase activity by treating cells with a small molecule that inhibits HDAC activity (TSA). We observed a loss of Kcnq1ot1-mediated silencing only at the genes it localizes to, but not neighboring genes (**Supplemental Figure 7B**). Together, these results demonstrate that Kcnq1ot1 localizes at a high concentration within the TAD containing its transcriptional locus, binds directly to SHARP, and recruits SHARP and its associated HDAC3 complex to silence transcription of genes within this nuclear compartment (**Figure 7H**).

In addition to Kcnq1ot1, we identified several other lncRNAs that demarcate specific nuclear compartments around their transcriptional loci containing their functional targets. For example: (**i)** Airn localizes within a TAD containing its reported imprinted target genes^110–112^ but excludes other neighboring genes in the genome (**Supplemental Figure 7D**,**G**). **(ii)** Pvt1 localizes to a TAD containing Myc and multiple known enhancers of Myc (**Supplemental Figure 7E**,**G**) and has been shown to repress Myc expression^113^. **(iii)** Chaserr localizes within the TAD containing Chd2 (**Supplemental Figure 7F**) and has been shown to repress Chd2 expression^114,115^.

These results demonstrate that the localization patterns of a ncRNA in 3D space guides the recruitment of regulatory proteins to these specific nuclear compartments and highlights an essential role for these RNA-mediated compartments in gene regulation.

## DISCUSSION

Our results demonstrate that RNAs localize within hundreds of compartments that occur throughout the entire nucleus, and that RNA is required for the localization of specific diffusible proteins (HP1, SHARP) and ncRNAs (e.g. U7, U1, snoRNAs, etc.) into dozens of compartmentalized structures. In all of these cases, we observed a common theme where (i) specific RNAs localize at high concentrations in spatial proximity to their transcriptional loci and (ii) diffusible ncRNA and protein molecules that bind to these RNAs are enriched within these compartmentalized structures. These observations suggest a common mechanism by which RNA can mediate nuclear compartmentalization: nuclear RNAs can form high concentration spatial territories close to their transcriptional loci (“seed”), bind to diffusible regulatory ncRNAs and proteins through high affinity interactions (“bind”) and by doing so, act to dynamically change the spatial distribution of these diffusible molecules in the nucleus such that they are enriched within compartments composed of multiple DNA loci, regulatory and target RNAs, and proteins in 3D space (“recruit”, **Figure 8**).

**Figure 8:**
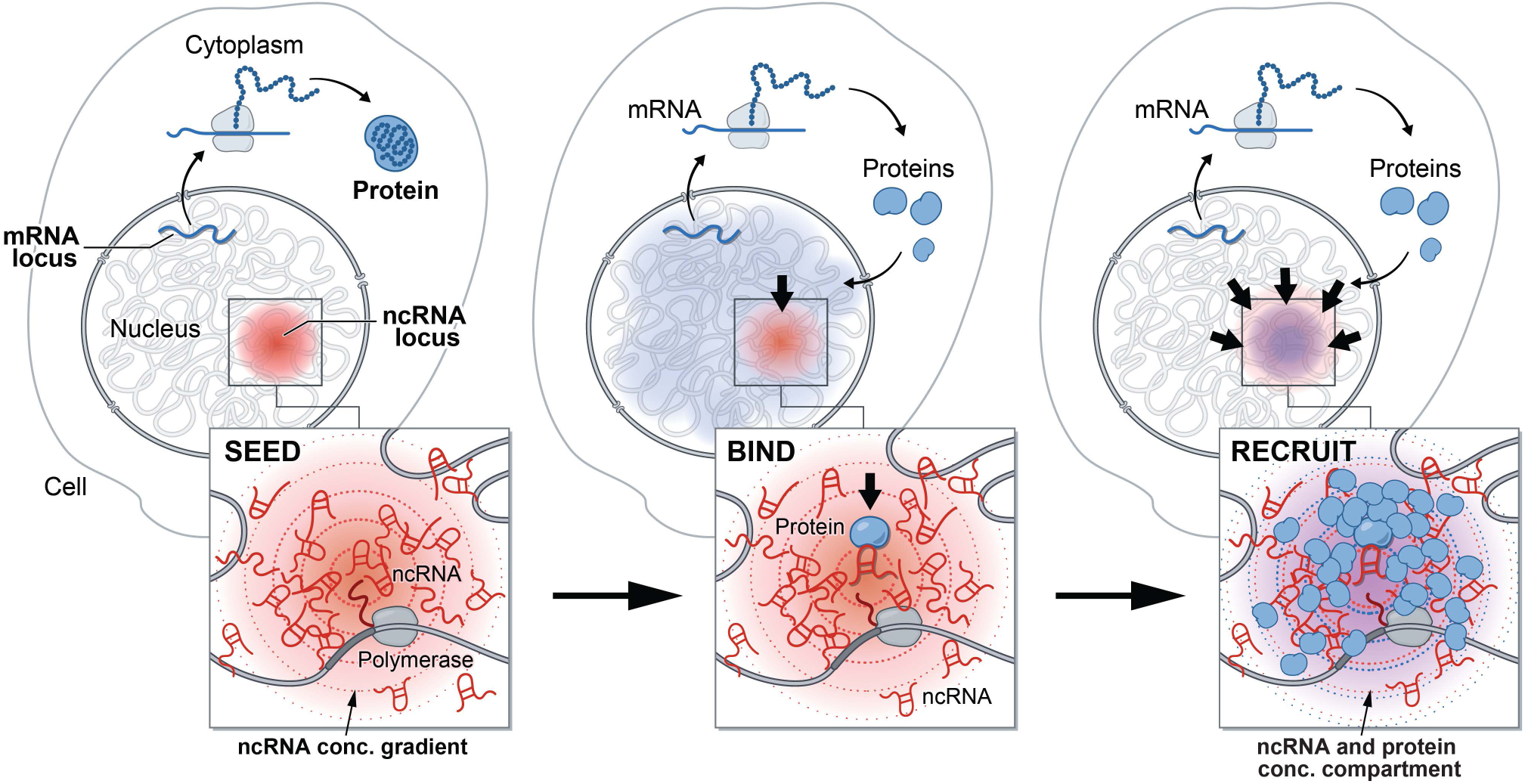
A model for the mechanism by which ncRNAs drive the formation of nuclear compartments. Upon transcription, mRNAs are exported to the cytoplasm (for translation to proteins) while ncRNAs are retained in the nucleus. The process of ncRNA transcription creates a concentration gradient of ncRNA transcript with the highest concentrations near its transcriptional locus (SEED, left panel). Because these RNAs are functional immediately upon transcription and can bind with high affinity to diffusible RNAs and proteins (BIND, middle panel), they can act to change the dynamic equilibrium of these proteins to concentrate them in a spatial compartment (RECRUIT, right panel). In this way, ncRNAs can drive the organization of regulatory and functional nuclear compartments containing RNA, DNA and proteins.

This mechanism may explain why many distinct types of RNA processing occur through compartmentalization of regulatory ncRNAs and proteins near their nascent RNA targets. Specifically, we show that each of these RNA processing hubs consists of a high concentration of nascent RNA near its transcriptional locus and enrichment of diffusible *trans*-associating ncRNAs – known to bind to the encoded nascent RNA – within the spatial compartment. In this way, these nuclear compartments contain high concentrations of regulatory RNAs and proteins in proximity to their nascent RNA targets, which are further organized within higher-order DNA structures that come together in 3D space to form distinct processing hubs. Because the efficiency of a biochemical reaction is increased when the substrate or enzyme concentration is increased, creating a high local concentration of regulators (e.g. spliceosomes) and targets (e.g. nascent pre-mRNAs) in 3D space may increase the kinetic efficiency of such reactions, and in turn increase the efficiency of co-transcriptional processing and regulation. This compartmentalization mechanism can also increase the rate at which regulators identify and engage targets, which may be particularly important in cases where the regulators (e.g. scaRNAs, U7) are expressed at low levels relative to their more abundant substrates (e.g. snRNAs, histone mRNAs). This spatial organization may be an important regulatory mechanism for ensuring the efficiency of co-transcriptional RNA processing and may explain how RNA processing and transcription are kinetically coupled.

Our results demonstrate that hundreds of nuclear ncRNAs are preferentially localized within precise structures in the nucleus, suggesting that this may be an important and common function exploited by additional nuclear RNAs to coordinate the spatial organization of diffusible molecules. This mechanism exploits a privileged and unique role for RNA in the nucleus (relative to DNA or proteins). Specifically, the process of transcription produces many copies of an RNA, which are by definition present at high concentrations in proximity to their transcriptional loci^18,116^. In contrast, proteins are translated in the cytoplasm and therefore lack positional information in the nucleus, and DNA is only present at a single copy and therefore cannot achieve high local concentrations.

Central to this mechanism is the fact that ncRNAs can form high affinity interactions with both protein and RNA immediately following transcription. In this way, they can act to recruit proteins and RNAs within these high concentration spatial compartments. In contrast, mRNAs are functional when translated into protein and do not form stable interactions with regulatory molecules in the nucleus. In this way, our results suggest that any RNA that functions independently of its translated product may similarly act as a ncRNA. For example, we note that nascent pre-mRNAs may also have protein-coding independent functions and form high-affinity interactions within the nucleus that are important for spatial organization. Indeed, we find that nascent pre-mRNAs and histone pre-mRNAs can seed organization of nuclear compartments even though their processed RNAs are also translated into protein products. This role for RNA as a seed for nuclear compartments might also explain formation of other recently described nuclear compartments such as transcriptional condensates^11,12^, which inherently produce high levels of RNA, including enhancer-associated RNAs and pre-mRNAs^117^. Nonetheless, not all ncRNAs – or even all nuclear ncRNAs – act to form compartments around their loci since nuclear ncRNAs can also localize within other regions in the nucleus (e.g. Malat1, scaRNAs, snoRNAs, and snRNAs). Future work will be needed to understand why some specific nuclear RNAs are constrained to local spatial compartments, while others diffuse throughout the nucleus.

This unique role for ncRNAs in the nucleus may explain why certain biological processes utilize ncRNA regulators rather than proteins or DNA. For example, coordinated regulation of multiple genomic DNA targets would be ideally controlled through the expression of a single ncRNA that could localize and recruit regulatory proteins to all of these targets simultaneously. Indeed, many multi-gene regulatory programs, such as X chromosome inactivation and imprinted gene silencing, utilize ncRNAs as regulators (e.g. Xist, Kcqn1ot1, and Airn). In this way, ncRNAs can increase both the efficiency and specificity of gene regulation by enabling control of multiple target genes through the expression of a single regulatory RNA from its genomic locus. This strategy may also be advantageous even when modulating a single gene because establishment of an RNA compartment can recruit effector proteins simultaneously to many genomic regions that are far away in linear distance but proximal in 3D space – including promoters and multiple enhancers – to enable higher concentration and more potent gene regulation. As an example, we observe high concentration of the Pvt1 lncRNA over the Myc gene and all of its known enhancer elements. This coordinated gene regulation model may extend to many of the hundreds of ncRNAs that we identified to be localized within discrete spatial compartments in the nucleus.

Taken together, these results provide a global picture of how spatial enrichment of ncRNAs in the nucleus can seed formation of compartments that coordinate the efficiency and specificity of a wide range of essential nuclear functions, including RNA processing, heterochromatin organization, and gene regulation (**Supplemental Figure 8**). While we focused our analysis on ncRNAs in this work, we note that RD-SPRITE can also be applied to measure how gene expression relates to genome organization because it can detect the arrangement of nascent pre-mRNAs relative other RNAs (e.g. enhancer RNAs, pre-mRNAs) and 3D DNA structure. Beyond the nucleus, we anticipate that RD-SPRITE will also provide a powerful method to study the molecular organization, function, and mechanisms of RNA compartments and granules throughout the cell.

## ACKNOWLEDGEMENTS

We thank Elizabeth Soehalim for help in adapting the SPRITE method; Sam Kim, Vickie Trinh, and Jasmine Thai for help with generating and validating cell lines and cell culture; Patrick McDonel for discussions and advice on the RD-SPRITE method and helpful comments on the manuscript; Andres Collazo for microscopy help; John Rinn, Drew Honson, Mackenzie Strehle, and Drew Perez for comments on the manuscript and helpful suggestions; Aaron Lin for sequencing help and advice; Shawna Hiley for editing; Inna-Marie Strazhnik and Sigrid Knemeyer for illustrations. S.A.Q. received support from the HHMI Gilliam Fellowship and NSF GRFP Fellowship. We thank Alexander Shishkin, Ward G. Walkup IV, Parham Peyda, and Vickie Trinh for help generating the Halo-tagged FL-SHARP vector. P.B. received support from NIH 5 T32 GM 7616-40, NIH NRSA CA247447, and the UCLA-Caltech Medical Scientist Training Program. N.O. is supported by the American Cancer Society Postdoctoral Fellowship (PF-17-240-01). J.W.J. is supported by a BBE post-doctoral fellowship from Caltech. A.K.B. was funded by NHLBI F30-HL136080 and the USC MD/PhD Program. Imaging was performed in the Biological Imaging Facility, with the support of the Caltech Beckman Institute and the Arnold and Mabel Beckman Foundation. This work was funded by the NIH 4DN (U01 DA040612 and U01 HL130007), the NYSCF, NIH Director’s Early Independence Award (DP5OD012190), CZI Ben Barres Early Career Acceleration Award, Sontag Foundation, Searle Scholars Program, Pew-Steward Scholars program, and funds from the California Institute of Technology. M.G. is a NYSCF-Robertson Investigator.

## AUTHOR CONTRIBUTIONS

S.A.Q. conceived of this project with M.G., led the development and optimization of the RD-SPRITE method, performed experiments, analyzed and interpreted data, generated figures, oversaw all aspects of the project, and wrote the paper. P.B. developed and optimized the RD-SPRITE protocol, performed SPRITE experiments, analyzed and interpreted data, contributed to data visualization, figure presentation, model schematics/illustrations, and wrote the paper. N.O. led the effort to analyze and interpret data, wrote software, created new methods for data analysis and visualization, performed analysis and visualization on the data and contributed major findings and results, created main and supplemental figures, and contributed to the initial draft of paper, model schematics/illustrations, and reviewed and edited the manuscript. J.W.J. designed, performed, acquired, and analyzed all the RNA-FISH, DNA-FISH, IF, IF/RNA-FISH experiments and made all imaging figures; performed all LNA-related experiments and generated the figures and results; performed Flavopiridol treatments and analysis; contributed to the writing of the centromeric RNA hub section, model schematics/illustrations, and provided comments and edits on the entire manuscript. A.K.B. performed all Kcnq1ot1 biochemical and functional experiments, including CRISPRi knockdowns, TSA treatments, and functional characterizations; worked with A.C. to develop and characterize the inducible Kcnq1ot1 cell line and to generate homozygous deletions of the SHARP Binding Site within Kcnq1ot1; worked with MRB to purify SHARP and map it to Kcnq1ot1. P.C. led the effort on the data processing and curation, writing scripts and constructing pipelines that enabled data interpretation; was responsible for gene, repeat, and allele annotation as well as validation and producing several QC metrics; contributed to experimental optimization of the RNA-DNA SPRITE protocol. M.R.B. developed the engineered SHARP lines for CLAP and methods for purification of SHARP; worked with A.K.B. to perform SHARP purifications for Kcnq1ot1 binding; advised and helped to develop and optimize the RNA molecular biology of the RD-SPRITE method in this project. A.C. developed all engineered cell lines used in this study, including the doxycycline inducible Xist cell lines, Kcnq1ot1 lines, SHARP binding site deletions, and dCas9 cell lines. Y.M. performed all live-cell 3D-SIM imaging and analysis of FL-SHARP and ΔRRM-SHARP localization. K.P. provided guidance and support on imaging, analysis, ideas, and discussions on the paper. M.G. conceived of this project with S.A.Q. and oversaw all experiments and analysis; performed computational analysis and generated scripts for analyzing the RD-SPRITE data; wrote the paper with S.A.Q. and P.B.

## MAIN FIGURE LEGENDS

**Supplemental Figure 1:**
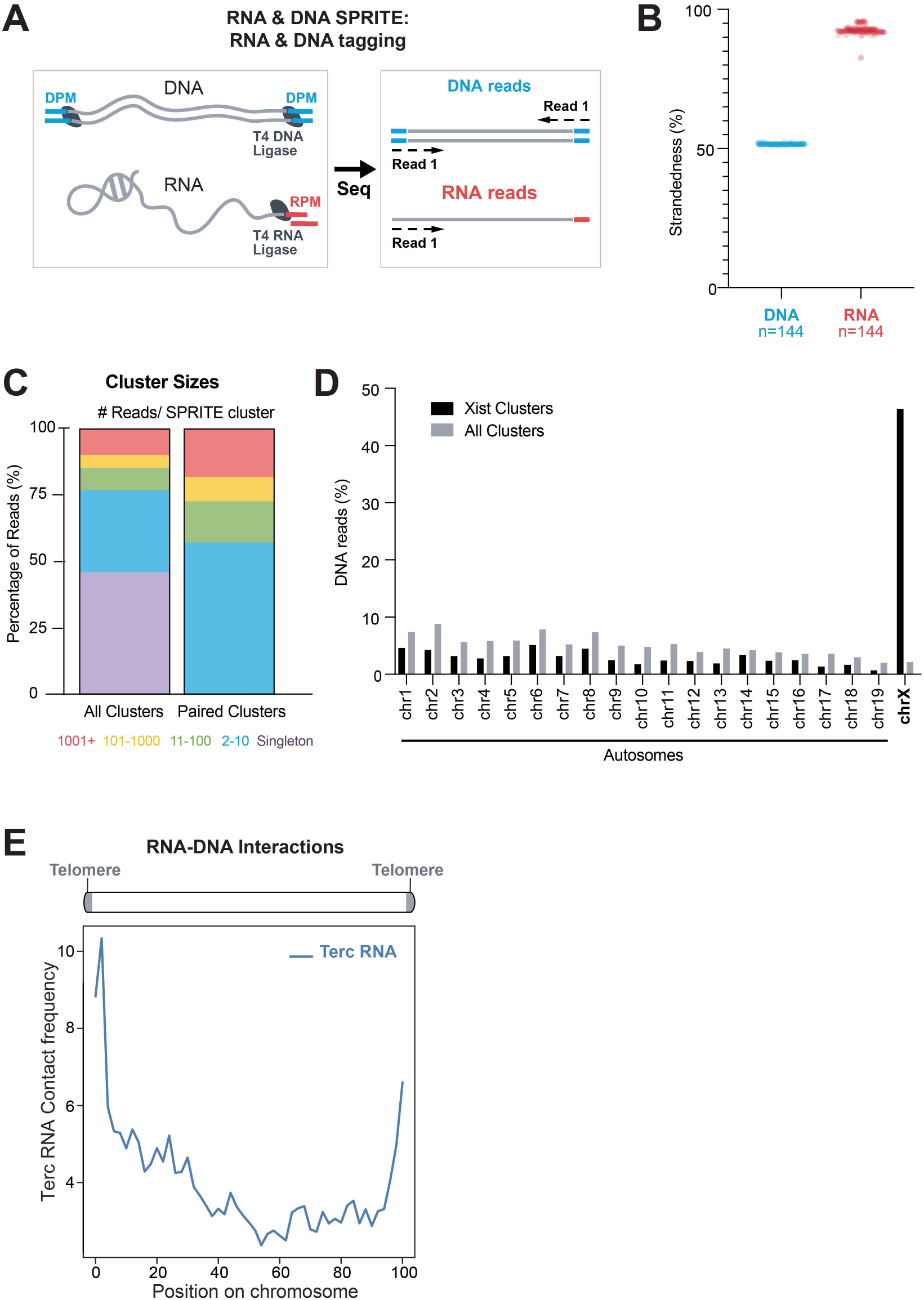
RD-SPRITE accurately measures RNA and DNA contacts. **(A)** Schematic of DNA and RNA tagging with sequence-specific tags to identify DNA- and RNA-specific reads through sequencing. DNA and RNA are each tagged with sequence-specific tags, namely “DNA Phosphate Modified” (DPM) tag and “RNA Phosphate Modified” (RPM) tags using T4 DNA and RNA Ligase, respectively. DNA is double stranded and therefore DPM will be read from both strands, while RNA is single stranded and therefore RPM will be read only from 1 strand. Additionally, the RPM and DPM tags have identical dsDNA sticky ends that enable subsequent split-pool barcoding with the same SPRITE tags. **(B)** The percentage of reads aligning to each DNA strand based on their DPM (DNA reads) or RPM (RNA reads) tags is shown across 144 independently amplified and sequenced SPRITE libraries from two SPRITE experiments. **(C)** Percentage of reads in SPRITE clusters of different sizes, stratified into categories of clusters containing 1, 2-10, 11-100, 101-1000, and 1001+ reads per cluster. Distributions shown for all clusters (left) and paired clusters (2+ reads per cluster) (right). **(D)** Percentage of DNA reads within each chromosome contained within SPRITE clusters containing the Xist RNA (black) compared to all SPRITE clusters (gray). **(E)** The aggregate unweighted contact frequency of the Telomerase associated RNA Component (Terc) across all chromosomes is shown.

**Supplemental Figure 2:**
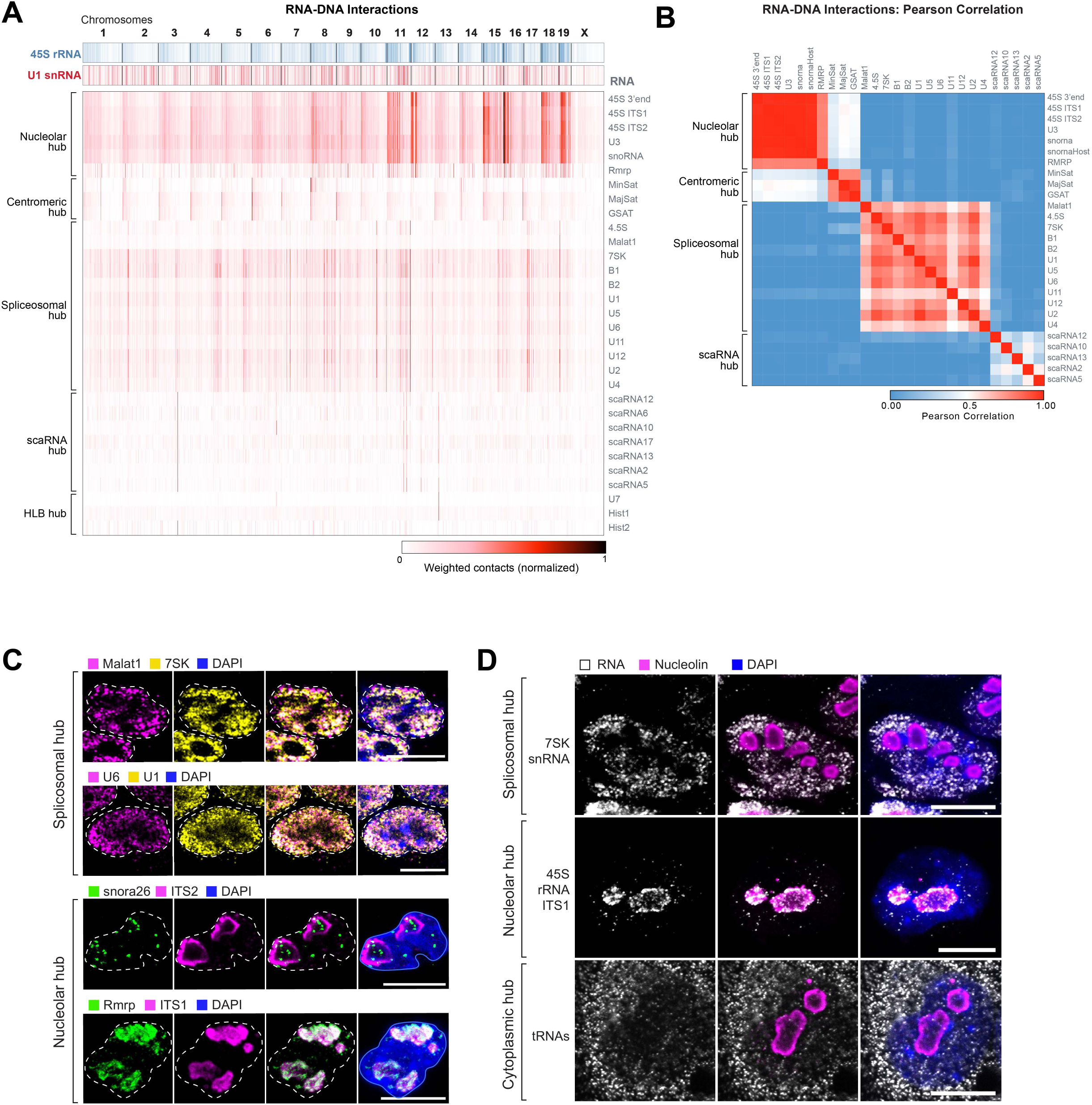
RNA localization on DNA and within the nucleus for RNAs within each RNA hub. **(A)** Weighted genomic DNA localization heatmap of each individual RNA. RNAs are organized by their RNA hub occupancy (shown in Figure 2A). Contacts are normalized from 0 to 1 to account for expression levels of each RNA. **(B)** Pearson correlation of RNA-DNA unweighted contact frequencies across the genome for individual RNAs within the nuclear hubs (nucleolar, centromeric, spliceosomal, and scaRNA hubs). Red represents high correlation and blue represents low correlation. **(C)** RNA FISH of various non-coding RNAs within the same hub in the nucleus. Spliceosomal hub (top): Malat1 lncRNA and 7SK RNA and (bottom): U6 and U1 spliceosomal RNAs. Nucleolar hub (top): snora26 snoRNA and 45S pre-rRNA ITS2 and (bottom): RNase MRP (Rmrp) and 45S pre-rRNA ITS1. Each panel is shown individually (left and middle) and overlaid (right). Dashed lines demarcate the nuclear boundary identified with DAPI. Scalebar is 10μm. **(D)** RNA FISH (left) along with nucleolin immunofluorescence (middle) and DAPI (right) of specific ncRNAs. 7SK RNA (top), ITS1 regions of 45S pre-rRNA (middle) and tRNAs (bottom). tRNAs are visualized using pooled RNA FISH probes (see Methods). Scalebar is 10μm.

**Supplemental Figure 3:**
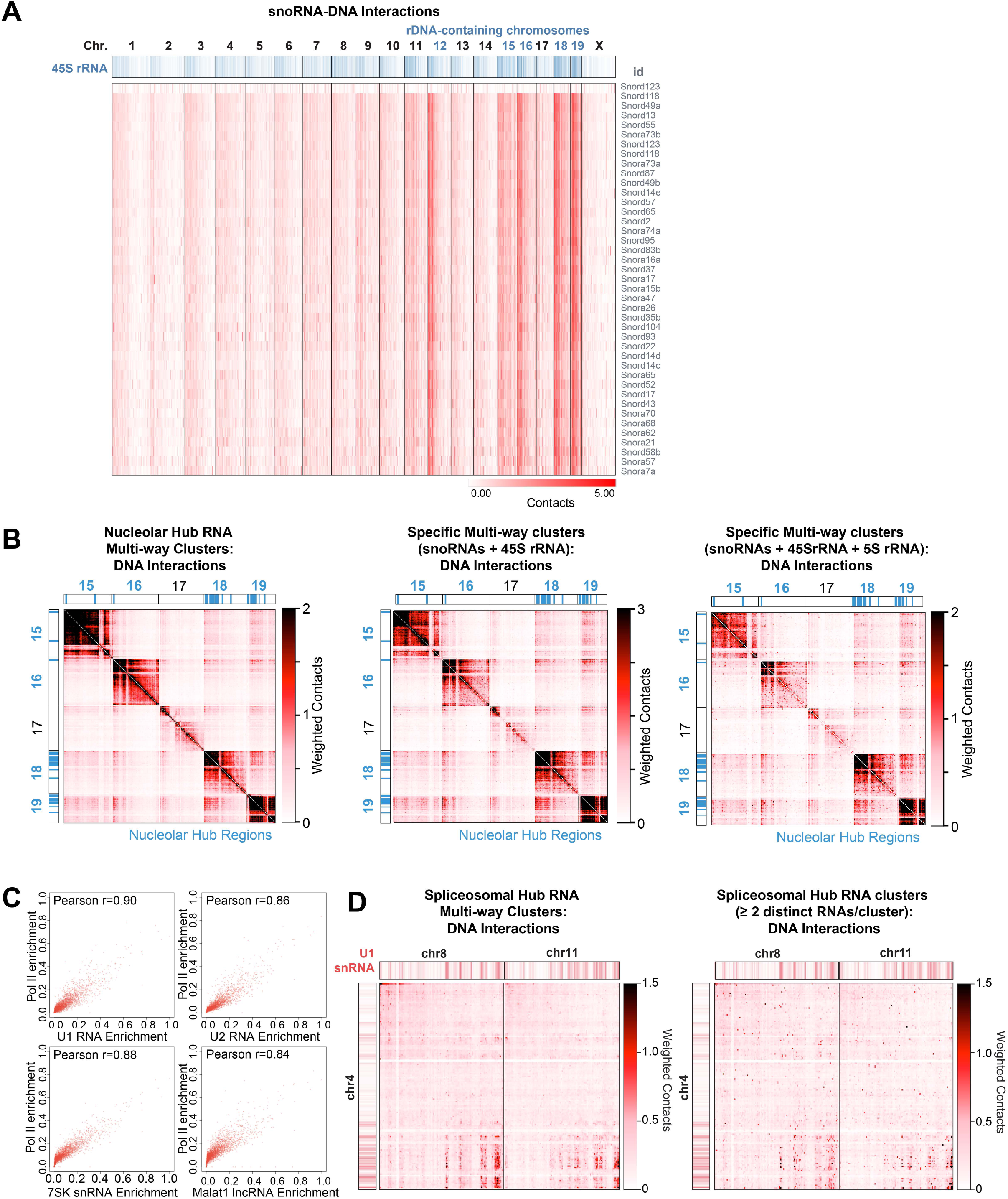
Nucleolar and spliceosomal hubs show higher-order interactions around loci of rRNA and mRNA genes, respectively. **(A)** Genome-wide localization of each individual snoRNA. Blue track shows 45S pre-rRNA localization on DNA. Chromosomes containing ribosomal DNA genes (chromosomes 12, 15, 16, 18, 19) are denoted in blue. (**B**) Weighted DNA-DNA contact heatmap shown for SPRITE clusters containing any of the RNAs within the nucleolar hub (left), both snoRNAs and 45S pre-rRNA (middle), and snoRNAs, 45S, and 5S (right) simultaneously. (**C**) 1Mb Enrichment of several spliceosomal hub RNA-DNA interactions (U1 snRNA, U2 snRNAs, 7SK RNA, and Malat1 lncRNA) compared to enrichment of Pol II ChIP-seq signal (ENCODE) genome wide. Pearson correlation provided for each set of RNA-DNA interactions and Pol II signal comparisons. (**D**) DNA-DNA contact heatmap shown for SPRITE clusters containing any of the RNAs contained within the spliceosomal hub (left) or containing 2 or more distinct spliceosomal hub RNAs simultaneously (right).

**Supplemental Figure 4:**
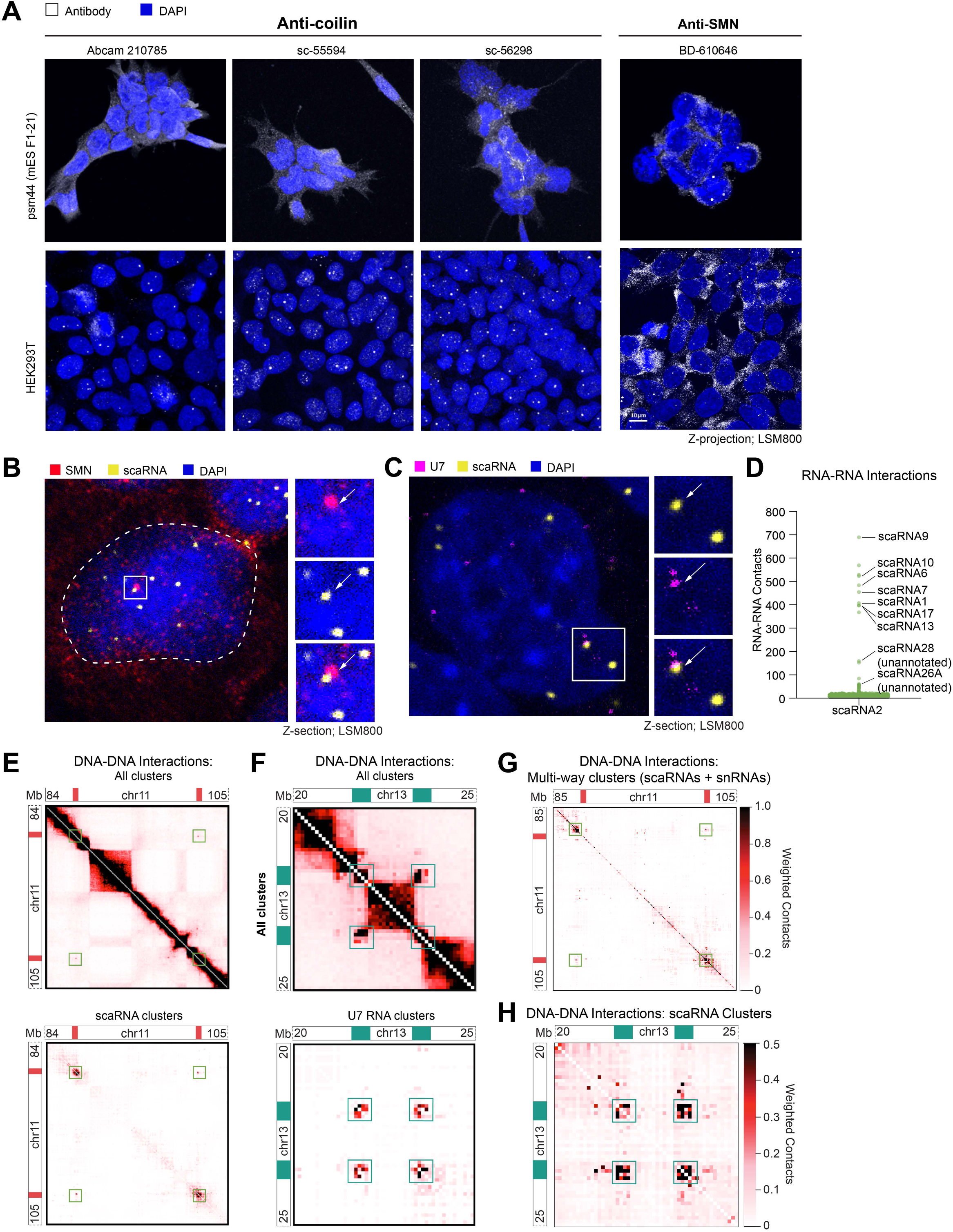
Spatial relationship between snRNA biogenesis hub and histone locus bodies. **(A)** Immunofluorescence imaging of classical Cajal Body (Coilin) and nuclear gem (SMN) markers in mouse ES cells and HEK293T cells. Top: Mouse ES cells do not contain visible Coilin foci for any of the three anti-Coilin antibodies tested. Bottom: HEK293T cells show visible Coilin foci. SMN foci, which are markers for nuclear Gemini of Cajal bodies (“gems”) are present in both mouse ES cells and HEK293T cells. **(B)** Z-section of mouse ES cell co-stained for SMN protein and scaRNAs (pooled scaRNA2 and scaRNA17 probes) within the nucleus (DAPI). Inset shows an example of scaRNA localization near SMN foci (arrow). **(C)** Z-section of mouse ES cell with RNA FISH staining for U7 and scaRNAs (pooled scaRNA2 and scaRNA17 probes) within the nucleus (DAPI). Inset shows an example of scaRNA localization near U7 (arrow). **(D)** RNA-RNA contact frequency between scaRNA2 and all RNAs. Top hits include annotated scaRNAs and identify two previously unannotated scaRNAs (see **Supplemental Methods**). **(E)** Weighted DNA-DNA contacts for all SPRITE clusters (top) and for SPRITE clusters containing scaRNAs (bottom) occurring within a region on chromosome 11 with snRNA gene clusters. scaRNA occupancy is demarcated with solid red boxes. **(F)** Weighted DNA-DNA contacts for all SPRITE clusters (top) and for SPRITE clusters containing the U7 ncRNA (bottom) occurring within a region on chromosome 13 containing the two Hist1 gene clusters. U7 and Hist1 RNA occupancy is demarcated with teal boxes. (**G**) Weighted DNA-DNA contacts shown for SPRITE clusters containing both scaRNAs and snRNAs simultaneously. **(H)** Weighted DNA-DNA contacts for SPRITE clusters containing the scaRNAs on chromosome 13.

**Supplemental Figure 5:**
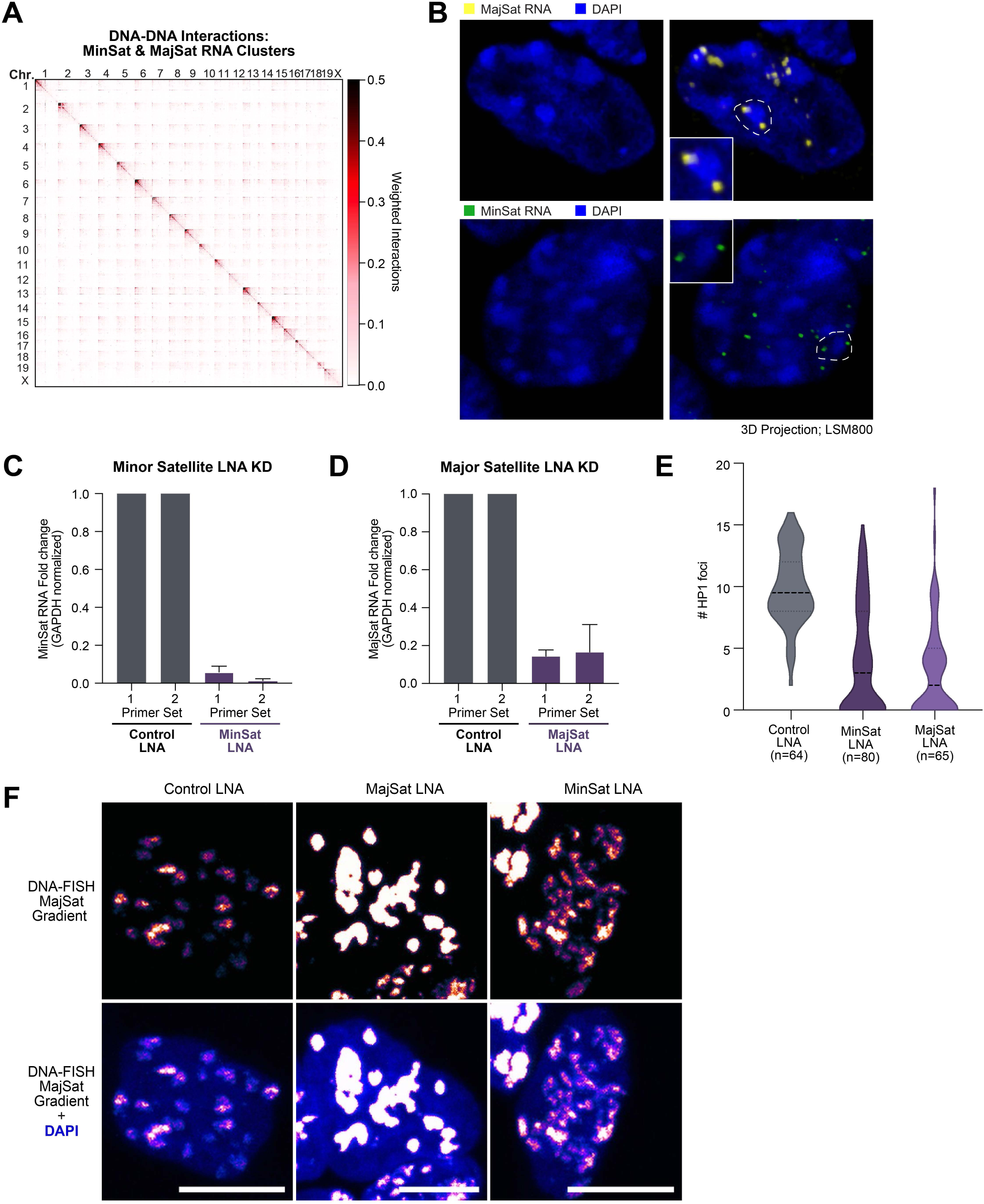
Satellite-derived ncRNAs mediate higher-order heterochromatin organization at centromeric clusters. **(A)** Weighted DNA-DNA contact matrices constructed from SPRITE clusters containing minor and major satellite RNAs. **(B)** 3D projections of either MajSat RNA FISH (top) or MinSat RNA FISH (bottom). DAPI in blue. Dashed lines and corresponding inset boxes zoom in on a single DAPI-dense chromocenter structure. **(C)** Quantification of mean LNA knockdown for minor satellite RNA (2 primer sets) compared to control LNA. Error bars represent standard deviation. **(D)** Quantification of LNA knockdown for major satellite RNA (2 primer sets) compared to control LNA. Error bars represent standard deviation. **(E)** Quantification of number of HP1 foci shown in Figure 5E (as a violin plot). Control: n=64 cells, MinSat: n=80 cells, MajSat: n=65 cells. **(F)** Imaging of DNA FISH in control (left) or LNA knockdown of MajSat (middle) and MinSat (right) RNA. DNA-FISH of MajSat shown as a color gradient indicating DNA FISH signal intensity (top) and with DAPI (bottom). See Supplemental Note 4 for detailed description of phenotypes. Scale bar is 10 μm.

**Supplemental Figure 6:**
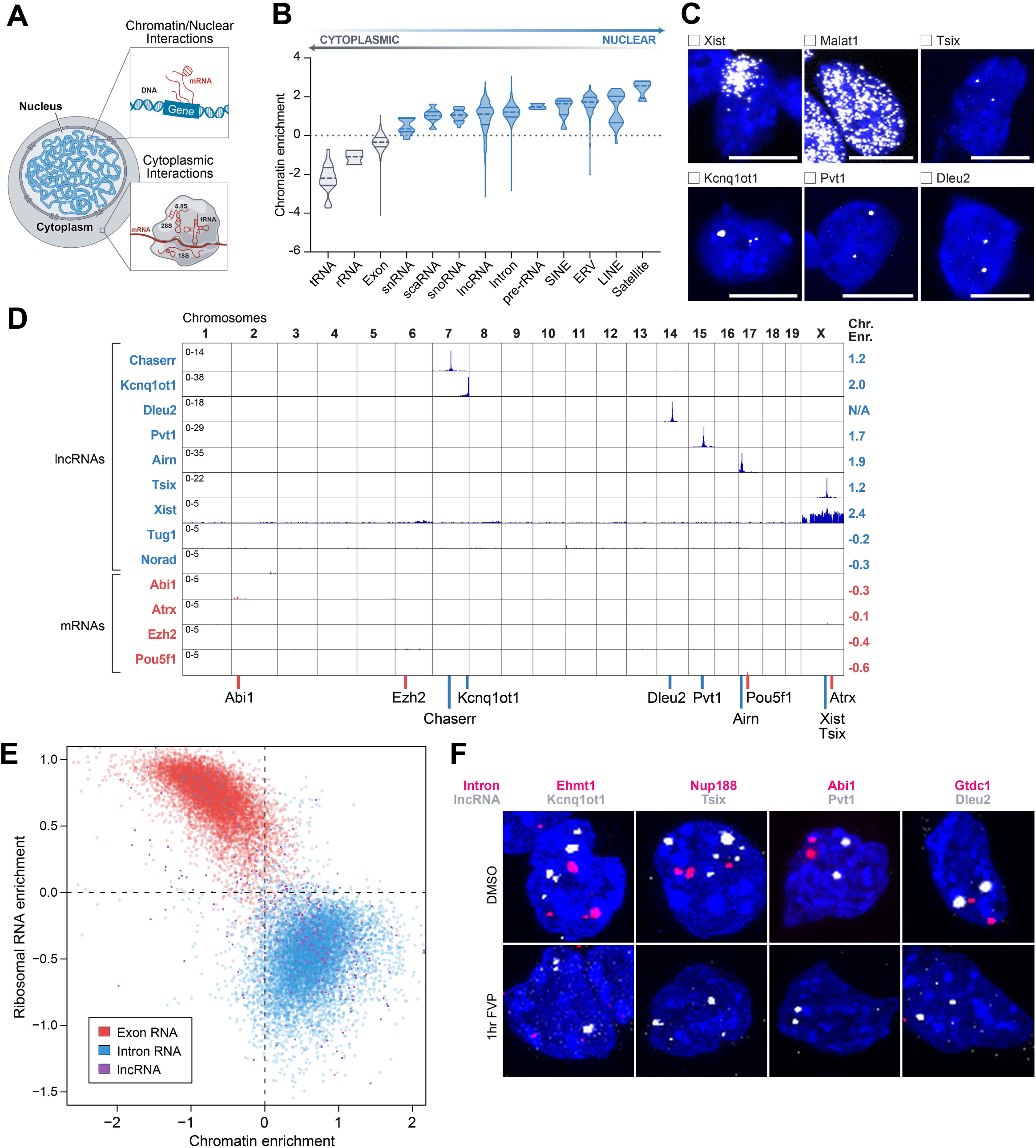
Many lncRNAs localize within 3D proximity to their transcriptional loci in the nucleus. **(A)** Schematic illustration of our chromatin enrichment score which computes the frequency of an RNA interaction with chromatin (top inset) compared to the frequency of interactions without chromatin, such as with rRNA, tRNA, and mRNA interactions in the cytoplasm (bottom inset). **(B)** Chromatin enrichment score for multiple classes of RNAs. tRNAs, rRNAs, and exons are predominantly depleted on chromatin (enrichment score < 0) versus other classes of RNAs, including introns, scaRNAs, lncRNAs, are enriched on chromatin (enrichment score > 0). **(C)** RNA FISH localization patterns of multiple lncRNAs (Xist, Malat1, Tsix, Kcnq1ot1, Pvt1, and Dleu2 lncRNAs) in the nucleus (DAPI). **(D)** Normalized RNA-DNA interactions for several lncRNAs (blue) and mRNAs (red). Each RNA locus is demarcated at the bottom. **(E)** Chromatin enrichment scores (x-axis) versus ribosomal RNA enrichment scores (y-axis) for exons (red), introns (blue), and lncRNAs (purple). **(F)** RNA FISH for 4 mRNA introns (Ehmt4, Nup188, Abi1, Gtdc2) and 4 lncRNAs (Kcnq1ot1, Tsix, Pvt1, Dleu2) treated for 1 hour with DMSO (top) or FVP (bottom). As a control, we co-stained lncRNAs (white) and introns (red) within the same cell.

**Supplemental Figure 7:**
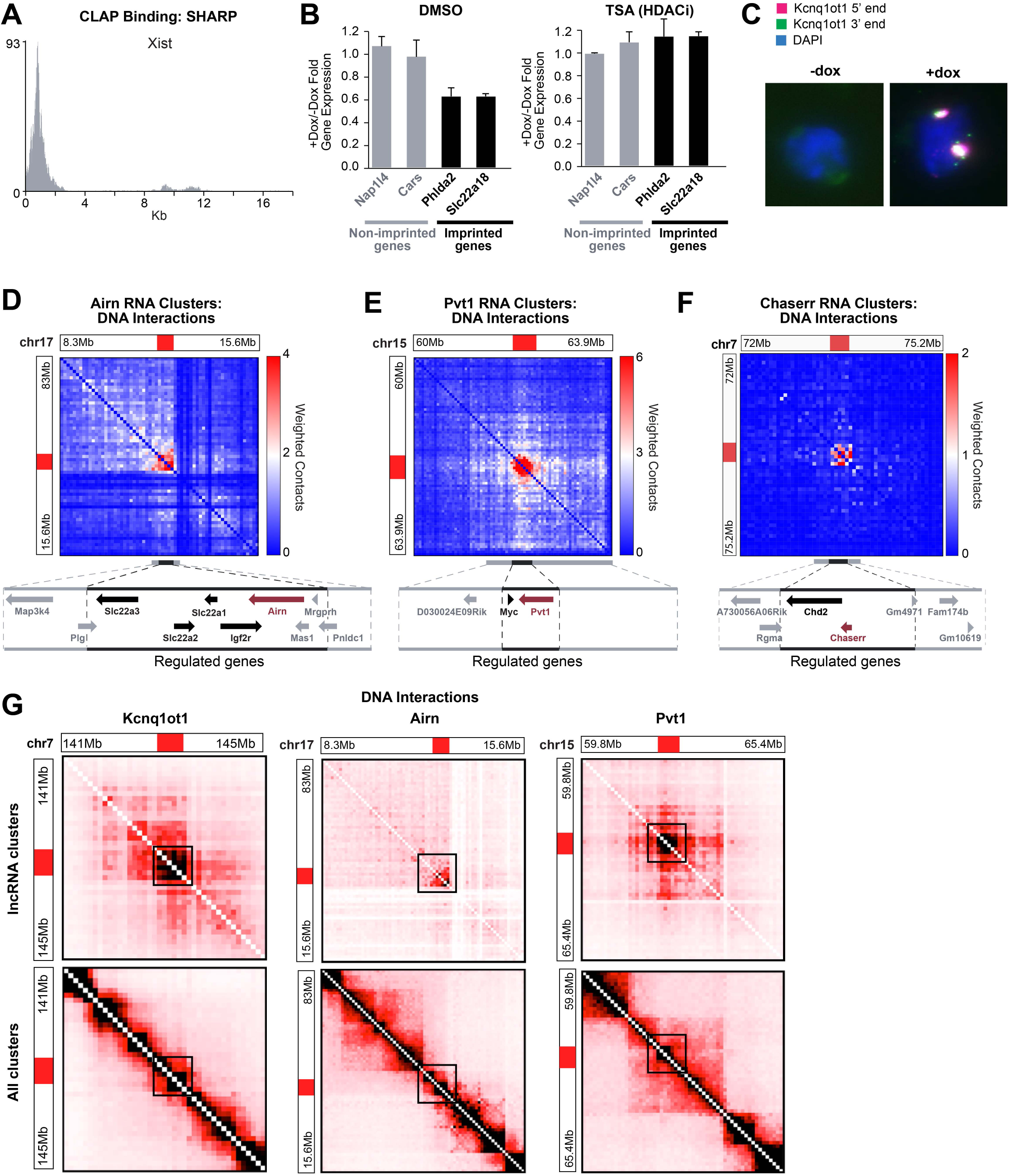
lncRNAs regulate target gene expression precisely within their localization domain. **(A)** SHARP protein binds to the Xist lncRNA, particularly at the 0-2kb region. **(B)** Mean gene expression differences of Kcnq1ot1-regulated and Kcnq1ot1-non-regulated genes between induced (+Dox) and non-induced (-Dox) samples treated with DMSO (left) or the HDAC inhibitor, Trichostatin A (TSA) (right). DMSO: Regulated genes show robust repression while genes not within the imprinted TAD show no difference in expression upon induction of Kcnq1ot1. TSA: Regulated genes show loss of repression and exhibit comparable gene expression changes with genes not within the imprinted TAD. Error bars represent standard deviation. **(C)** RNA FISH performed with two distinct probes targeting the Kcnq1ot1 lncRNA – 1 probe set was designed against the 3’ end of the RNA and the other designed against the 5’ end of the RNA. FISH was performed in cells in the absence of doxycycline (left) and in the presence of doxycycline (right). **(D)** Weighted DNA-DNA interaction matrix for Pvt1 RNA-containing SPRITE clusters showing Pvt1 lncRNA localization on DNA in a region occupied by Pvt1 and Myc genes. **(E)** Weighted DNA-DNA interaction matrix for Airn RNA-containing SPRITE clusters showing Airn lncRNA localization on DNA in a region confined to the genes Airn is known to regulate^118^. **(F)** Weighted DNA-DNA interaction matrix for Chaserr RNA-containing SPRITE clusters. Chaserr RNA is confined to a TAD containing the Chaserr gene and its known regulatory target, Chd2. **(G)** Top: DNA-DNA contacts within SPRITE clusters containing each lncRNA (left: Kcnq1ot1, middle: Airn, right: Pvt1) and bottom: DNA-DNA contacts across all SPRITE clusters within the same regions.

**Supplemental Figure 8:**
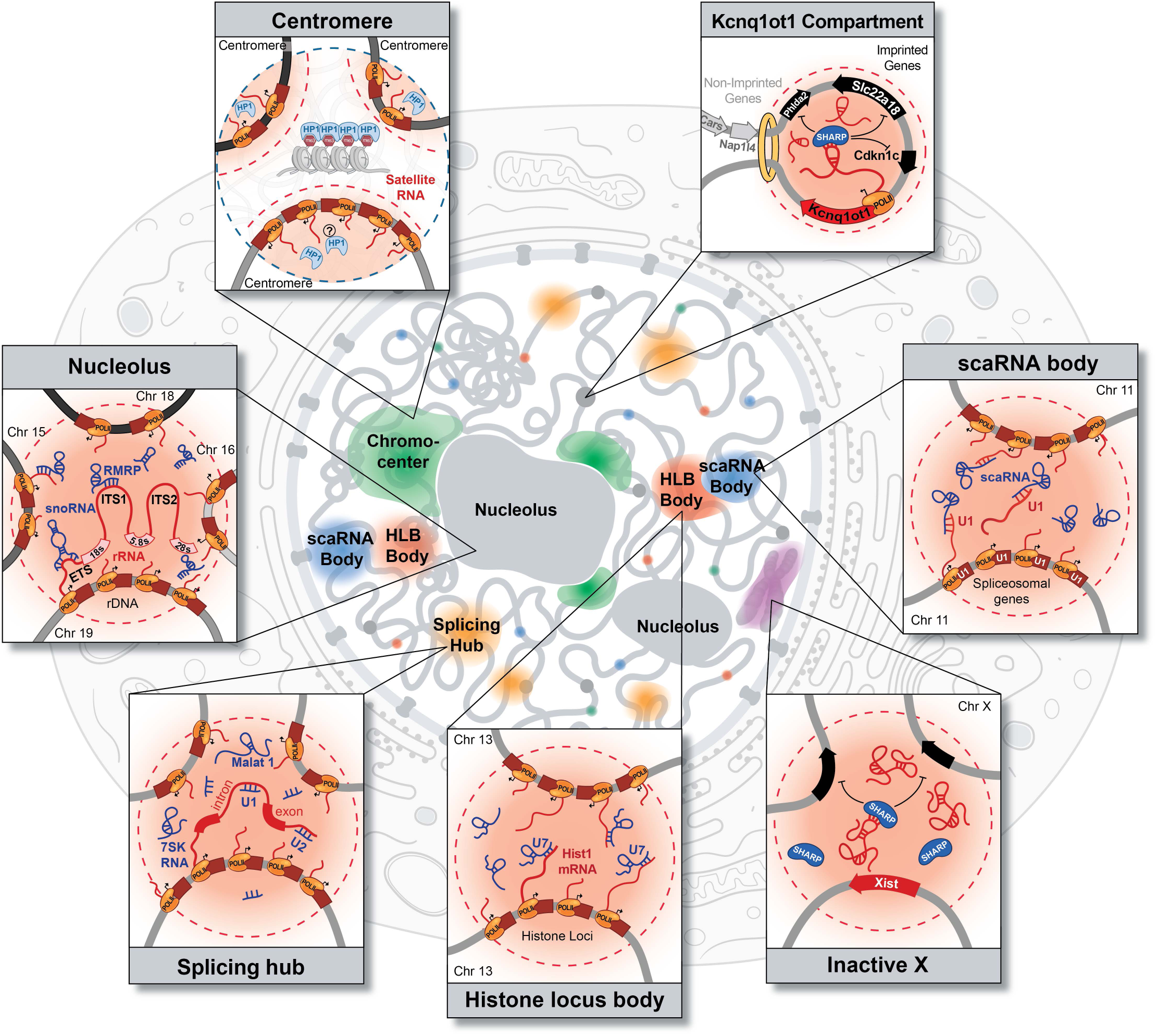
A widespread role for ncRNAs in shaping compartments throughout the nucleus that are associated with various nuclear functions. A schematic of the localization of the different nuclear compartments within the nucleus and the molecular components contained within them. In each of these cases, an RNA seeds organization by achieving high concentration in spatial proximity to its transcriptional locus. This leads to the formation of nuclear compartments associated with RNA processing, heterochromatin assembly, and gene regulation.

## SUPPLEMENTAL FIGURE LEGENDS

**Supplemental Video 1**: **Full length SHARP localizes in discrete diffraction-limited foci**. Live-cell 3D-SIM of Halo-tagged FL-SHARP JF646 captured for ∼2 minutes reveals distinct and persistent SPEN foci throughout the nucleus.

**Supplemental Video 2**: **Deletion of the RNA recognition motifs of SHARP leads to diffusive localization**. Live-cell 3D-SIM of ΔRRM-SHARP JF646 captured for ∼2 minutes exhibits a diffusive localization pattern and no observable foci in the nucleus.

## SUPPLEMENTAL NOTES

**Supplemental Note 1: *RD-SPRITE improves efficiency of RNA tagging***. Although our previous version of SPRITE could map both RNA and DNA, it was limited primarily to detecting highly abundant RNA species (e.g. 45S pre-rRNA). In RD-SPRITE, we have improved detection of lower abundance RNAs by increasing yield through the following adaptations. (i) We increased the RNA ligation efficiency by utilizing a higher concentration of RPM, corresponding to ∼2000 molar excess during RNA ligation. (ii) Adaptor dimers that are formed through residual purification on our magnetic beads lead to reduced efficiency because they preferentially amplify and preclude amplification of tagged RNAs. To reduce the number of adaptor dimers in library generation, we introduced an exonuclease digestion of excess reverse transcription (RT) primer that dramatically reduces the presence of the RT primer. (iii) Reverse transcription is used to add the barcode to the RNA molecule, yet when RT is performed on crosslinked material it will not efficiently reverse transcribe the entire RNA (because crosslinked proteins will act to sterically preclude RT). To address this, we performed a short RT in crosslinked samples followed by a second RT reaction after reverse crosslinking to copy the remainder of the RNA fragment. (iv) Because cDNA is single stranded, we need to ligate a second adaptor to enable PCR amplification. The efficiency of this reaction is critical for ensuring that we detect each RNA molecule. We significantly improved cDNA ligation efficiency by introducing a modified “splint” ligation. Specifically, a double stranded “splint” adaptor containing the Read1 Illumina priming region and a random 6mer overhang is ligated to the 3’end of the cDNA at high efficiency by performing a double stranded DNA ligation. This process is more efficient than the single stranded DNA-DNA ligation previously utilized^54^. (v) Finally, we found that nucleic acid purification performed after reverse crosslinking leads to major loss of complexity because we lose a percentage of the unique molecules during each cleanup. In the initial RNA-DNA SPRITE protocol there were several column (or bead) purifications utilized to remove enzymes and enable the next enzymatic reaction. We reduced these cleanups by introducing biotin modifications into the DPM and RPM adaptors that enable binding to streptavidin beads and for all subsequent molecular biology steps to occur on the same beads. Together, these improvements enabled a dramatic improvement of our overall RNA recovery and enables generation of high complexity RNA/DNA structure maps.

**Supplemental Note 2: *The snRNA biogenesis hub may be similar to the Cajal body***. We note that the snRNA biogenesis hub may be similar to Cajal bodies, which have been noted to contain snRNA genes and scaRNAs^82,84,86,119,120^. However, Cajal bodies are traditionally defined by the presence of Coilin foci in the nucleus^80,84,121^ and based on this definition, our mES cells do not contain visible Cajal bodies with all three antibodies tested (**Supplemental Figure S4A**). Despite the absence of traditionally defined Cajal bodies, our data suggest that snRNA biogenesis hubs do indeed exist and form around snRNA gene loci, even in the absence of observable Coilin foci. Our data suggest that scaRNA localization more accurately defines snRNA processing bodies relative to Coilin. Consistent with this idea, scaRNAs have a clearly defined functional role in snRNA biogenesis whereas Coilin is dispensable for snRNA biogenesis^86^. It is also possible that these snRNA processing bodies are distinct from Cajal bodies, which may represent a different nuclear structure. For example, these might represent nuclear gems^122^, which contain SMN protein, or “residual bodies,” which are Coilin negative^123,124^. We note that we observe SMN foci in our mES cells and that some, but not all, scaRNAs colocalize with SMN protein in the nucleus (**Supplemental Figure S4A-B**).

**Supplemental Note 3: *RD-SPRITE measures the frequency at which RNAs are contacting chromatin***. Although data from previous methods have reported that both lncRNAs and mRNAs are similarly enriched on chromatin at their transcriptional loci, we observed a striking difference in chromatin localization between these classes of RNA. The major reason for this is because RD-SPRITE measures RNA localization within all compartments of the cell, including in the nucleus and cytoplasm. Accordingly, we can compute a chromatin enrichment score, which we define as the frequency at which a given RNA is localized on chromatin (**Supplemental Figure 6A-B**). Other RNA-DNA mapping methods such as hybridization (e.g. RAP, ChIRP) or proximity-ligation (e.g. GRID-Seq, Margi) methods exclusively measure RNA when they are present on chromatin and therefore cannot measure this differential localization frequency.

**Supplemental Note 4: *Depletion of satellite-derived ncRNAs leads to altered chromocenter structure***. Chromocenter regions display altered structure 48h and 72h post MajSat and MinSat LNA transfection in comparison to control. All 3 conditions (MajSat, MinSat, Control) were visualized by DNA-FISH (chromocenters clustering depicted in gradient of fluorescent intensity from MajSat targeting probes) and DAPI staining (chromocenters depicted as DAPI rich regions). The observed phenotypes include: (i) larger pericentromeric chromatin foci and clusters in comparison to control samples; (ii) smaller pericentromeric chromatin clusters and foci that seem to be less condensed (rod-shaped) in comparison to controls (**Supplemental Figure 5F**). Our results demonstrate that both MajSat and MinSat RNAs are required for HP1*β* recruitment to chromocenter regions.

## MATERIALS AND METHODS

### Lead Contact

Further information and requests for resources and reagents should be directed to and will be fulfilled by the Lead Contact, Mitchell Guttman.

### Materials Availability

This study did not generate new unique reagents.

### Data and Code Availability

The analysis pipeline used in this study is available at https://github.com/GuttmanLab/sprite2.0-pipeline. Datasets generated during this study will be available on GEO.

### Cell line generation, cell culture, and drug treatments

#### Cell lines used in this study

We used the following cell lines in this study: (i) Female ES cells (*pSM44* ES cell line) derived from a 129 × castaneous F1 mouse cross. These cells express Xist from the endogenous locus under control of a tetracycline-inducible promoter. The dox-inducible Xist gene is present on the 129 allele, enabling allele-specific analysis of Xist induction and X chromosome silencing. (ii) Female ES cells where we replaced the endogenous Kcnq1ot1 promoter with a tetracycline-inducible promoter on both alleles (*Kcnq1ot1-inducible* ES cell line). In the absence of Doxycycline, these cells do not express Kcnq1ot1 from either allele; in the presence of Doxycycline, these cells express Kcnq1ot1 biallelically. (iii) Female ES cells containing dCas9 fused to 4-copies of the SID transcriptional repression domain integrated into a single locus in the genome (dCas9-4XSID). (iv) HEK293T, a female human embryonic kidney cell line obtained from ATCC.

#### Cell culture conditions

All mouse ES cell lines were cultured in serum-free 2i/LIF medium as previously described^54^. HEK293T cells were cultured in complete media consisting of DMEM (GIBCO, Life Technologies) supplemented with 10% FBS (Seradigm Premium Grade HI FBS, VWR), 1X penicillin-streptomycin (GIBCO, Life Technologies), 1X MEM non-essential amino acids (GIBCO, Life Technologies), 1 mM sodium pyruvate (GIBCO, Life Technologies) and maintained at 37°C under 5% CO_2_. For maintenance, 800,000 cells were seeded into 10 mL of complete media every 3-4 days in 10 cm dishes. HEK293T cells were used for human-mouse mixing experiments to assess noise during the SPRITE procedure as well as for imaging Coilin foci.

#### Doxycycline Inducible Cell Line Development

Female ES cells (F1 2-1 line, provided by K. Plath) were CRISPR-targeted (nicking gRNA pairs TGGGCGGGAGTCTTCTGGGCAGG and GGATTCTCCCAGGCCCAGGGCGG) to integrate the Tet transactivator (M2rtTA) into the Rosa26 locus using R26P-M2rtTA, a gift from Rudolf Jaenisch (Addgene plasmid #47381). This line was subsequently CRISPR-targeted (nicking gRNA pairs GCTCGTTTCCCGTGGATGTG and GCACGCCTTTAACTGATCCG) to replace the endogenous Xist promoter with tetracycline response elements (TRE) and a minimal CMV promoter as previously described^46^. The promoter replacement insertion was verified by PCR amplification of the insertion locus and Sanger sequencing of the amplicon. SNPs within the amplicon allowed for allele identification of the insertion, confirming that the 129 allele was targeted and induced Xist expression. We routinely confirmed the presence of two X chromosomes within these cells by checking the presence of X-linked SNPs on the 129 and castaneous alleles.

#### 3D-SIM SHARP-Halo cell culture conditions

pSM33 cells were seeded in 4-well imaging chambers (ibidi) equipped with a high precision glass bottom and plasmids were transfected with lipofectamine 3000 24 hours prior to imaging according to the manufacturer’s instructions. Addition of doxycycline 8hrs prior to imaging was performed to induce SHARP expression. 1μM JF646 Halo ligand was introduced to the media for 30 min, washed-off twice with PBS and exchanged with fresh media which were incubated for another 15 min. Live-cell 3D-SIM imaging was performed at 37C and 5% CO_2_ in media without phenol red.

#### Doxycycline Inducible Kcnq1ot1 lines

The endogenous promoter of Kcnq1ot1 was CRISPR-targeted (nicking gRNA pairs TCGTGGCTGCCACGTCACCA and CAGATGCTGAATAATGACTA) to insert a TRE and minimal CMV promoter. Clones were screened for ablation of endogenous Kcnq1ot1 expression and biallelic upregulation of expression upon administration of doxycycline using RNA FISH probes targeting the 5’end and 3’end of the RNA (**Supplemental Figure 7C**).

#### CRISPRi: dCas9-4XSID cell line generation

A catalytically dead Cas9 (dCas9) fused to 4 copies of the SID repressive domain (4XSID) expressed from an Ef1*α* promoter was integrated into a single copy locus in the genome (mm10 - chr6:86,565,487-86,565,506; gRNA sequence AATCTTAGTACTACTGCTGC) using CRISPR targeting (cells hereby referred to as dCas9-4XSID).

#### Doxycycline induction

Xist and Kcnq1ot1 expression were induced in their respective cell lines by treating cells with 2 μg/ml doxycycline (Sigma). Xist was induced for 24 hours prior to crosslinking and analysis. Kcnq1ot1 was induced for 12-16hrs prior to RNA harvesting.

#### Trichostatin (TSA) treatment

For HDAC inhibitor experiments, cells were treated with either DMSO (control) or 5μM TSA (Sigma T8552-1MG) in fresh 2i media or 2μg/ml doxycycline in standard 2i.

#### Flavopiridol (FVP) Treatment

FVP transcriptional inhibition was performed by culturing cells in FVP (Sigma F3055-1MG) or DMSO at 1uM final concentration for 1 hour.

### RNA & DNA-SPRITE: Simultaneous tagging of RNA and DNA interactions using SPRITE

RD-SPRITE is an adaptation of our initial SPRITE protocol^54^ with significant improvements to the RNA molecular biology steps that enable generation of higher complexity RNA libraries. The approach was performed as follows:

#### Crosslinking, lysis, sonication, and chromatin digestion

Cells were lifted using trypsinization and were crosslinked in suspension at room temperature with 2mM disuccinimidyl glutarate (DSG) for 45 minutes followed by 3% Formaldehyde for 10 minutes to preserve RNA and DNA interactions *in situ*. After crosslinking, the formaldehyde crosslinker was quenched with addition of 2.5M Glycine for final concentration of 0.5M for 5 minutes, cells were spun down, and resuspended in 1x PBS + 0.5% RNAse Free BSA (AmericanBio #AB01243-00050) over three washes, 1x PBS + 0.5% RNAse Free BSA was removed, and flash frozen at −80C for storage. We found that RNAse Free BSA is critical to avoid RNA degradation. RNase Inhibitor (1:40, NEB Murine RNAse Inhibitor or Thermofisher Ribolock) was also added to all lysis buffers and subsequent steps to avoid RNA degradation. After lysis, cells were sonicated at 4-5W of power for 1 minute (pulses 0.7 second on, 3.3 seconds off) using the Branson Sonicator and chromatin was fragmented using DNAse digestion to obtain DNA of approximately ∼150bp-1kb in length.

#### Estimating molarity

After DNase digestion, crosslinks were reversed on approximately 10 μl of lysate in 82 μL of 1X Proteinase K Buffer (20 mM Tris pH 7.5, 100 mM NaCl, 10 mM EDTA, 10 mM EGTA, 0.5% Triton-X, 0.2% SDS) with 8 μL Proteinase K (NEB) at 65°C for 1 hour. RNA and DNA were purified using Zymo RNA Clean and Concentrate columns per the manufacturer’s specifications (>17nt protocol) with minor adaptations, such as binding twice to the column with 2X volume RNA Binding Buffer combined with by 1X volume 100% EtOH to improve yield. Molarities of the RNA and DNA were calculated by measuring the RNA and DNA concentration using the Qubit Fluorometer (HS RNA kit, HS dsDNA kit) and the average RNA and DNA sizes were estimated using the RNA High Sensitivity Tapestation and Agilent Bioanalyzer (High Sensitivity DNA kit).

#### NHS bead coupling

We used the RNA and DNA molarity estimated in the lysate to calculate the total number of RNA and DNA molecules per microliter of lysate. We coupled the lysate to NHS-activated magnetic beads (Pierce) in 1x PBS + 0.1% SDS combined with 1:40 dilution of NEB Murine RNase Inhibitor overnight at 4°C as previously described^54^. We coupled at a ratio of 0.5 molecules per bead to reduce the probability of simultaneously coupling multiple independent complexes to the same bead, which would lead to their association during the split-pool barcoding process. Because multiple molecules of DNA and RNA can be crosslinked in a single complex, this estimate is a more conservative estimate of the number of molecules to avoid collisions on individual beads. After NHS coupling overnight, the coupling was quenched in 0.5M Tris pH 7.5 and beads were washed post coupling as previously described.

Because the crosslinked complexes are immobilized on NHS magnetic beads, we can perform several enzymatic steps by adding buffers and enzymes directly to the beads and performing rapid buffer exchange between each step on a magnet. All enzymatic steps were performed with shaking at 1200 rpm (Eppendorf Thermomixer) to avoid bead settling and aggregation. All enzymatic steps were inactivated either by adding 1 mL of SPRITE Wash buffer (20mM Tris-HCl pH 7.5, 50mM NaCl, 0.2% Triton-X, 0.2% NP-40, 0.2% Sodium deoxycholate) supplemented with 50 mM EDTA and 50 mM EGTA to the NHS beads or Modified RLT buffer (1x Buffer RLT supplied by Qiagen, 10mM Tris-HCl pH 7.5, 1mM EDTA, 1mM EGTA, 0.2% N-Lauroylsarcosine, 0.1% Triton-X, 0.1% NP-40).

#### DNA End Repair and dA-tailing

We then repair the DNA ends to enable ligation of tags to each molecule. Specifically, we blunt end and phosphorylate the 5′ ends of double-stranded DNA using two enzymes. First, T4 Polynucleotide Kinase (NEB) treatment is performed at 37°C for 1 hour, the enzyme is quenched using 1 mL Modified RLT buffer, and then buffer is exchanged with two washes of 1 mL SPRITE Detergent Buffer to beads at room temperature. Next, the NEBNext End Repair Enzyme cocktail (containing T4 DNA Polymerase and T4 PNK) and 1x NEBNext End Repair Reaction Buffer is added to beads and incubated at 20°C for 1 hour, and inactivated and buffer exchanged as specified above. DNA was then dA-tailed using the Klenow fragment (5′-3′ exo-, NEBNext dA-tailing Module) at 37°C for 1 hour, and inactivated and buffer exchanged as specified above. Note, we do not use the combined NEB End Repair/dA tailing modules as the temperatures in the protocol are not compatible with SPRITE as the higher temperature will reverse crosslinks. To prevent degradation of RNA, each enzymatic step is performed with the addition of 1:40 NEB Murine RNAse Inhibitor or Thermofisher Ribolock.

#### Ligation of the DNA Phosphate Modified (“DPM”) Tag

After end repair and dA-tailing of DNA, we performed a pooled ligation with “DNA Phosphate Modified” (DPM) tag that contains certain modifications that we found to be critical for the success of RD-SPRITE. Specifically, (i) we incorporate a phosphothiorate modification into the DPM adaptor to prevent its enzymatic digestion by Exo1 in subsequent RNA steps and (ii) we integrated an internal biotin modification to facilitate an on-bead library preparation post reverse-crosslinking. The DPM adaptor also contains a 5’phosphorylated sticky end overhang to ligate tags during split-pool barcoding. Ligation was performed as previously described using Instant Sticky End Mastermix (NEB) except that all ligations were supplemented with 1:40 RNAse inhibitor (ThermoFisher Ribolock or NEB Murine RNase Inhibitor) to prevent RNA degradation. Because T4 DNA Ligase only ligates to double-stranded DNA, the unique DPM sequence enables accurate identification of DNA molecules after sequencing.

#### Ligation of the RNA Phosphate Modified (“RPM”) Tag

To map RNA and DNA interactions simultaneously, we ligated a RNA adaptor to RNA that contains the same 7nt 5’phosphorylated sticky end overhang as the DPM adaptor to ligate tags to both RNA and DNA during split-pool barcoding. To do this, we first modify the 3’end of RNA to ensure that they all have a 3’OH that is compatible for ligation. Specifically, RNA overhangs are repaired with T4 Polynucleoide Kinase (NEB) with no ATP at 37°C for 20 min. RNA is subsequently ligated with a “RNA Phosphate Modified” (RPM) adaptor as previously described using High Concentration T4 RNA Ligase I^125^. Because T4 RNA Ligase 1 only ligates to single-stranded RNA, the unique RPM sequence enables accurate identification of RNA and DNA molecules after sequencing. After RPM ligation, RNA was converted to cDNA using Superscript III at 42°C for 1 hour using the “RPM bottom” RT primer that contains an internal biotin to facilitate on-bead library construction (as above) and a 5’end sticky end to ligate tags during SPRITE. Excess primer is digested with Exonuclease 1. All ligations were supplemented with 1:40 RNAse inhibitor (ThermoFisher Ribolock or NEB Murine RNase Inhibitor) to prevent RNA degradation.

#### Split-and-pool barcoding to identify RNA and DNA interactions

The beads were then repeatedly split- and-pool ligated over four rounds with a set of “Odd,” “Even” and “Terminal” tags (see SPRITE Tag Design in Quinodoz et al. Cell 2018^54^). Both DPM and RPM contain the same 7 nucleotide sticky end that will ligate to all subsequent split-pool barcoding rounds. All split-pool ligation steps and reverse crosslinking were performed for 45min to 1 hour at 20°C as previously described. All ligations were supplemented with 1:40 RNAse inhibitor (ThermoFisher Ribolock or NEB Murine RNase Inhibitor) to prevent RNA degradation.

#### Reverse crosslinking

After multiple rounds of SPRITE split-and-pool barcoding, the tagged RNA and DNA molecules are eluted from NHS beads by reverse crosslinking overnight (∼12-13 hours) at 50°C in NLS Elution Buffer (20mM Tris-HCl pH 7.5, 10mM EDTA, 2% N-Lauroylsarcosine, 50mM NaCl) with added 5M NaCl to 288mM NaCl Final combined with 5uL Proteinase K (NEB).

#### Post reverse-crosslinking library preparation

AEBSF (Gold Biotechnology CAS#30827-99-7) is added to the Proteinase K (NEB Proteinase K #P8107S; ProK) reactions to inactive the ProK prior to coupling to streptavidin beads. Biotinylated barcoded RNA and DNA are bound to streptavidin beads. To improve recovery, the supernatant is bound again to 20 ul of streptavidin beads and combined with the first capture. Beads are washed in 1X PBS + RNase inhibitor and then resuspended in 1x First Strand buffer to prevent any melting of the RNA:cDNA hybrid. Beads were pre-incubated at 40C for 2 min to prevent any sticky barcodes from annealing. A second reverse transcription is performed by adding Superscript III (without RT primer) to extend the cDNA through the areas which were previously crosslinked. The second RT ensures that cDNA recovery is maximal, particularly if RT terminated at a crosslinked site prior to reverse crosslinking. After generating cDNA, the RNA is degraded by addition of RNaseH and RNase cocktail, and the 3’end of the resulting cDNA is ligated to attach an dsDNA oligo containing library amplification sequences for subsequent amplification.

Previously, we performed cDNA (ssDNA) to ssDNA primer ligation which relies on the two single stranded sequences coming together for conversion to a product that can then be amplified for library preparation. To improve the efficiency of cDNA molecules ligated with the Read1 Illumina priming sequence, we perform a “splint” ligation, which involves a chimeric ssDNA-dsDNA adaptor that contains a random 6mer that anneals to the 3’ end of the cDNA and brings the 5’ phosphorylated end of the cDNA adapter directly together with the cDNA via annealing. This ligation is performed with 1x NEB Instant Sticky End Master Mix at 20°C for 1 hour. This greatly improves the cDNA tagging and overall RNA yield.

Libraries were amplified using Q5 Hot-Start Mastermix (NEB) with primers that add the full Illumina adaptor sequences. After amplification, the libraries are cleaned up using 0.8X SPRI (AMPure XP) and then gel cut using the Zymo Gel Extraction Kit selecting for sizes between 280 bp - 1.3 kb.

#### Sequencing

Sequencing was performed on an Illumina NovaSeq S4 paired-end 150×150 cycle run. For the mES RNA-DNA RD-SPRITE data in this experiment, 144 different SPRITE libraries were generated from two technical replicate SPRITE experiments and were sequenced. Each SPRITE library corresponds to a distinct aliquot during the Proteinase K reverse crosslinking step which is separately amplified with a different barcoded primer, providing an additional round of SPRITE barcoding.

#### Primers Used for RPM, DPM, and Splint Ligation (IDT)

1. RPM top: /5Phos/rArUrCrArGrCrACTTAGCG TCAG/3SpC3/
2. RPM bottom (internal biotin): /5Phos/TGACTTGC/iBiodT/GACGCTAAGTGCTGAT
3. DPM Phosphorothioate top: /5Phos/AAGACCACCAGATCGGAAGAGCGTCGTG*T* A*G*G* /32MOErG/ *Denotes Phosphorothioate bonds
4. DPM bottom (internal biotin): /5Phos/TGACTTGTCATGTCT/iBioT/CCGATCTGGTGGTCTT
5. 2Puni splint top: TACACGACGCTCTTCCGATCT NNNNNN/3SpC3/
6. 2Puni splint bottom: /5Phos/AGA TCG GAA GAG CGT CGT GTA/3SpC3/

#### Annealing of adaptors

A double-stranded DPM oligo and 2P universal “splint” oligo were generated by annealing the complementary top and bottom strands at equimolar concentrations. Specifically, the oligos were annealed in 1x Annealing Buffer (0.2 M LiCl_2_, 10 mM Tris-HCl pH 7.5) by heating to 95°C and then slowly cooling to room temperature (−1°C every 10 sec) using a thermocycler.

#### Assessing molecule to bead ratio

We ensured that SPRITE clusters represent *bona fide* interactions that occur within a cell by mixing human and mouse cells and ensuring that virtually all SPRITE clusters (∼99%) represent molecules exclusively from a single species. Specifically, we separately crosslinked HEK293T cells performed a human-mouse mixing RD-SPRITE experiment and identified conditions with low interspecies mixing (molecules = RNA+DNA instead of DNA). Specifically, for SPRITE clusters containing 2-1000 reads, the percent of interspecies contacts is: 2 beads:molecule = 0.9% interspecies contacts, 4 beads:molecule = 1.1% interspecies contacts, 8 beads:molecule = 1.1% interspecies contacts. We used the 2 beads:molecule ratio for the RD-SPRITE data set generated in this paper.

### RD-SPRITE processing pipeline

#### Adapter trimming

Adapters were trimmed from raw paired-end fastq files using Trim Galore! v0.6.2 (https://www.bioinformatics.babraham.ac.uk/projects/trim_galore/) and assessed with Fastqc v0.11.9. Subsequently, the DPM (GATCGGAAGAG) and RPM (ATCAGCACTTA) sequences are trimmed using Cutadapt v2.5^126^ from the 5’ end of Read 1 along with the 3’ end DPM sequences that result from short reads being read through into the barcode (GGTGGTCTTT, GCCTCTTGTT, CCAGGTATTT, TAAGAGAGTT, TTCTCCTCTT, ACCCTCGATT). The additional trimming helps improve read mapping in the end-to-end alignment mode. The SPRITE barcodes of trimmed reads are identified with Barcode ID v1.2.0 (https://github.com/GuttmanLab/sprite2.0-pipeline) and the ligation efficiency is assessed. Reads with an RPM or a DPM barcode are split into two separate files, to process RNA and DNA reads individually downstream, respectively.

#### Processing RNA reads

RNA reads were aligned to GRCm38.p6 with the Ensembl GRCm38 v95 gene model annotation using Hisat2 v2.1.0^127^ with a high penalty for soft-clipping --sp 1000,1000. Unmapped and reads with a low MapQ score (samtools view -bq 20) were filtered out for downstream realignment. Mapped reads were annotated for gene exons and introns with the featureCounts tool from the subread package v1.6.4 using Ensembl GRCm38 v95 gene model annotation and the Repeat and Transposable element annotation from the Hammel lab^128^. Filtered reads were subsequently realigned to our custom collection of repeat sequences using Bowtie v2.3.5^129^, only keeping mapped and primary alignment reads.

#### Processing DNA reads

DNA reads were aligned to GRCm38.p6 using Bowtie2 v2.3.5, filtering out unmapped and reads with a low MapQ score (samtools view -bq 20). Data generated in F1 hybrid cells (pSM33: C57BL/6 × 129SV-Jae or pSM44: 129 × castaneous) were assigned the allele of origin using SNPsplit v0.3.4^130^. RepeatMasker^131^ defined regions with milliDev <= 140 along with blacklisted v2 regions were filtered out using Bedtools v2.29.0^132^.

#### SPRITE cluster file generation

RNA and DNA reads were merged, and a cluster file was generated for all downstream analysis. MultiQC v1.6^133^ was used to aggregate all reports.

#### Masked bins

In addition to known repeat containing bins, we manually masked the following bins (mm10 genomic regions: chr2:79490000-79500000, chr11:3119270-3192250, chr15:99734977-99736026, chr3:5173978-5175025, chr13:58176952-58178051) because we observed a major overrepresentation of reads in the input samples.

### Microscopy imaging

#### 3D-Structured Illumination Microscopy (3D-SIM)

3D-SIM super-resolution imaging was performed on a DeltaVision OMX-SR system (Cytiva, Marlborough, MA, USA) equipped with a 60x/1.42 NA Plan Apo oil immersion objective (Olympus, Tokyo, Japan), sCMOS cameras (PCO, Kelheim, Germany) and 642 nm diode laser. Image stacks were acquired with z-steps of 125 nm and with 15 raw images per plane. The raw data were computationally reconstructed with the soft-WoRx 7.0.0 software package (Cytiva, Marlborough, MA, USA) using a wiener filter set to 0.002 and channel-specifically measured optical transfer functions (OTFs) using an immersion oil with a 1.518 refractive index (RI). 32-bit raw datasets were imported to ImageJ and converted to 16-bit stacks.

#### Immunofluorescence (IF)

Cells were grown on coverslips and rinsed with 1xPBS, fixed in 4% paraformaldehyde in PBS for 15 minutes at room temperature, rinsed in 1xPBS, and permeabilized with 0.5% Triton X-100 in PBS for 10 minutes at room temperature. Cells were either stored at −20°C in 70% ethanol or used directly for immunostaining and incubated in blocking solution (0.2% BSA in PBS) for at least 1 hour. If stored in 70% ethanol, cells were re-hydrated prior to staining by washing 3 times in 1xPBS and incubated in blocking solution (0.2% BSA in PBS) for at least 1 hour. Primary antibodies were diluted in blocking solution (see below) and added to coverslips for 3-5 hours at room temperature incubation. Cells were washed three times with 0.01% Triton X-100 in PBS for 5 minutes each and then incubated in blocking solution containing corresponding secondary antibodies labeled with Alexa fluorophores (Invitrogen) for 1 hour at room temperature. Next, cells were washed 3 times in 1xPBS for 5 minutes at room temperature and mounting was done in ProLong Gold with DAPI (Invitrogen, P36935). Images were collected on a LSM800 confocal microscope (Zeiss) with a 63× oil objective. *Z* sections were taken every 0.3 μm. Image visualization and analysis was performed with Icy software and ImageJ software.

#### Antibodies

Primary antibodies used in the study: anti-Nucleolin (Abcam ab22758 1:500); anti-NPAT (Abcam ab70595, 1:100); anti-SMN (BD 610646, 1:100); anti-CENP-A (Cell Signaling C51A7, 1:500); anti-HP1beta (Active Motif 39979, 1:200); anti-Coilin (Abcam Ab-210785, Santa Cruz sc-55594, Santa Cruz sc-56298, 1:100), all diluted in blocking solution.

#### RNA Fluorescence in situ Hybridization (RNA-FISH)

RNA-FISH performed in this study was based on the ViewRNA ISH (Thermo Fisher Scientific, QVC0001) protocol with minor modifications. Cells grown on coverslips were rinsed in 1xPBS, fixed in 4% paraformaldehyde in 1xPBS for 15 minutes at room temperature, permeabilized in 0.5% Triton-100 in the fixative for 10 minutes at room temperature, rinsed 3 times with 1xPBS and stored at −20°C in 70% ethanol until hybridization steps. All the following steps were performed according to manufacturer’s recommendations. Coverslips were mounted with ProLong Gold with DAPI (Invitrogen, P36935) and stored at 4°C until acquisition. For nuclear and nucleolar RNAs, cells were pre-extracted with 0.5% ice cold Triton-100 for 3 minutes to remove cytoplasmic background and fixed as described. All probes used in the study were custom made by Thermofisher. To test their specificity, we either utilized RNAse treatment prior to RNA-FISH or two different probes targeting the same RNA. Images were acquired on Zeiss LSM800 confocal microscope with a 100x glycerol immersion objective lens and Z-sections were taken every 0.3 μm. Image visualization and analysis was performed with Icy software and ImageJ software.

RNA FISH for scaRNA and tRNAs were performed with a combined set of probes to increase the signal of lower abundance RNAs. Specifically, scaRNAs were visualized with two combined probes of scaRNA2 and scaRNA17. tRNAs were visualized using probes targeting tRNA-Arg-TCG-4-1, tRNA-Leu-AAG-3-1, tRNA-Ile-AAT-1-8, tRNA-Arg-TCT-5-1, tRNA-Leu-CAA-2-1, tRNA-Ile-TAT-2-1, tRNA-Tyr-GTA-1-1. tRNA sequences were obtained using the GtRNAdb GRCm38/mm10 predictions (Lowe Lab, UCSC)^134,135^.

#### RNA-FISH and IF

For immunostaining combined with in situ RNA visualization, we used the ViewRNA Cell Plus (Thermo Fisher Scientific, 88-19000-99) kit per the manufacturer’s protocol with minor modifications. First immunostaining was performed as described above but all the incubations were performed in blocking buffer with addition of RNAse inhibitor and all the wash steps were done in RNAse free 1xPBS with RNAse inhibitor. Blocking buffer, PBS, RNAse inhibitors are provided in a kit. After the last wash in 1xPBS, cells underwent post-fixation in 2% paraformaldehyde on 1xPBS for 10min at room temperature, were washed 3 times in 1XPBS, and then RNA-FISH protocol was followed as described above. Images were acquired on the Zeiss LSM800 confocal microscope with a 100x glycerol immersion objective lens and Z-sections were taken every 0.3 μm. Image visualization and analysis was performed with Icy software (http://icy.bioimageanalysis.org/) and ImageJ software (https://imagej.nih.gov/).

#### RNA-FISH for FVP experiments

To compare the relative stability of lncRNAs and pre-mRNAs, we obtained intron FISH probes for targets of comparable gene length to lncRNAs. This was done to ensure that any differences in RNA stability upon FVP treatment are not due to differences in the time it takes to transcribe each RNA. Specifically, we obtained probes for pre-mRNAs that are 57.87kb (Nup188), 73.7kb (Mbd5), 99.8kb (Abi1), 129.7kb (Ehmt1),131.8kb (Atrx), and 297.2kb (Gtdc1) in length. For lncRNAs, we obtained probes for RNAs of lengths 53.4kb (Tsix), 79.5kb (Dleu2), 93.1kb (Kcnq1ot1), and 340kb (Pvt1).

#### DNA-FISH

DNA-FISH was performed as previously described^136^ with modifications. Cells grown on coverslips were rinsed with 1xPBS, fixed in 4% paraformaldehyde in 1xPBS for 15 minutes at room temperature, permeabilized in 0.5% Triton-100 in the fixative for 10 minutes at room temperature, rinsed 3 times with 1xPBS and stored at −20°C in 70% ethanol until hybridization steps. Pre-hybridization cells were dehydrated in 100% ethanol and dried for 5 minutes at room temperature. 4ul drop of hybridization mix with probes was spotted on a glass slide and dried coverslips were placed on the drop. Coverslips were sealed with rubber cement, slides were incubated for 5 minutes at 85°C, and then incubated overnight at 37°C in humid atmosphere. After hybridization and three washes with 2xSSC, 0.05% Triton-100 and 1mg/ml PVP in PBS at 50°C for 10 minutes, cells were rinsed in 1xPBS and mounted with ProLong Gold with DAPI (Invitrogen, P36935).

Hybridization buffer consisted of 50% formamide, 10% dextran sulphate, 2xSSC, 1 mg/ml polyvinyl pyrrolidone (PVP), 0.05% Triton X-100, 0.5 mg/ml BSA. 1 mM short oligonucleotides labeled with Cy5 ([CY5]ttttctcgccatattccaggtc) were used as probes against Major Satellites and full-length minor satellite repeat sequence was used as probes against Minor Satellites. Minor satellite sequence was firstly cloned to pGEM plasmid and then labeled by PCR reaction with self-made TAMRA dATPs for minor satellites. Labeled PCR product was purified with a QIAquick PCR Purification Kit (QIAGEN) and 50ng was mixed with hybridization buffer. Images were acquired on Zeiss LSM800 confocal microscope with a 63x glycerol immersion objective lens and Z-sections were taken every 0.3 μm. Image visualization and analysis was performed with Icy software and ImageJ software.

### Analysis of RNA-DNA contacts

#### Generating contact profiles

To map the genome-wide localization profile of a specific RNA, we calculated the contact frequency between the RNA transcript and each region of the genome binned at various resolutions (1Mb, 100kb and 10kb). Raw contact frequencies were computed by counting the number of SPRITE clusters in which an RNA transcript and a genomic bin co-occur. We normalized these raw contacts by weighting each contact by a scaling factor based on the size of its corresponding SPRITE cluster. Specifically, we enumerate all pairwise contacts within a SPRITE cluster and weight each contact by 2/n, where n is the total number of reads within a cluster.

#### RNA and cluster sizes

RNA-DNA contacts were computed for a range of SPRITE cluster sizes, such as 2-10, 11-100, and 101-1000 reads. We found that different RNAs tend to be most represented in different clusters sizes – likely reflecting the size of the nuclear compartment that they occupy. For example, 45S and snoRNAs are most represented in large clusters, while Malat1, snRNAs, and other ncRNAs tend to be represented in smaller SPRITE clusters. For analyses in this paper we utilized clusters containing 2-1000 reads.

#### Visualizing contact profiles

These methods produce a one-dimensional vector of DNA contact frequencies for each RNA transcript that we output in bedgraph format and visualize with IGV^137^. To compare DNA contact profiles between RNA transcripts, we calculated a Pearson correlation coefficient between the one-dimensional DNA contact vectors for all pairs of RNA transcripts.

#### Aggregate analysis

To map RNA localization across chromosomes with respect to centromeres and telomeres (e.g. Terc and satellite ncRNAs), we computed an average localization profile as a function of distance from the centromere of each chromosomes. To do this, we converted each 1Mb genomic bin into a percentile bin from 0 to 100 based on its relative position on its chromosome (from 5’ to 3’ ends). We then calculated the average contact frequency between a given RNA and each percentile bin across all chromosomes.

#### Allele specific analysis

To map localization to different alleles, we identified all clusters containing a given RNA (as above) and quantified the number of DNA reads uniquely mapping to each allele using allele specific alignments. Allele specific RNA-DNA contact frequencies were normalized by overall genomic read coverage for each allele to account for differences in coverage for each allele.

#### Nucleolar hub RNA-DNA contacts

We observe enrichment of pre-rRNAs and other nucleolar hub RNAs on chromosomes containing 45S ribosomal DNA (rDNA). Specifically, rDNA genes are contained on the centromere-proximal regions of chromosomes 12, 15, 16, 18, and 19 in mouse ES cells. We previously showed that regions on these chromosomes organize around nucleoli in the majority of cells imaged with DNA FISH combined with immunofluorescence for Nucleolin^54^. We also observed nucleolar hub RNAs enriched on other genomic regions corresponding to centromere-proximal DNA and transcriptionally inactive, gene poor regions. We previously showed that these genomic regions are organized proximal to the nucleolus using SPRITE and microscopy^54^.

### Analysis of RNA-RNA contacts

#### RNA-RNA contact matrices

We computed contact frequency between each RNA-RNA pair by counting the number of SPRITE clusters containing two different RNAs. To account for coverage differences in individual RNAs, we normalized this matrix using a matrix balancing normalization approach as previously described^138^. Briefly, this approach works by ensuring the rows and columns of a symmetric matrix add up to 1. In this way, RNA abundance does not dominate the overall strength of the contact matrix. For multi-copy RNAs (e.g. repeat-encoded RNAs, ribosomal RNA, tRNAs), all reads mapping to a given RNA were collapsed. Specifically, multi-copy RNA reads mapping to either the mm10 genome annotated using repeat masker or a custom repeat genome consensus were collapsed.

#### RNA Hubs

RNAs in each hub were identified using hierarchical clustering of the RNA-RNA contact matrix. Specifically, each hub corresponds to sets of RNAs with high contacts with other RNAs within the same hub, but low contacts with other RNAs in other hubs.

#### Mapping intron versus exon RNA-RNA contacts

To explore the differential RNA contacts that occur within nascent pre-mRNA and mature mRNAs, we focused on the intronic regions and exonic regions of mRNAs respectively. We retained all intronic or exonic regions that were contained in at least 100 independent SPRITE clusters. We then generate contact matrices between splicing non-coding RNAs (U1, U2, U4, U5, U6) and translation non-coding RNAs (18S, 28S, 5S, 5.8S) and these mRNA exons, and introns. We performed a matrix balancing normalization (ICE normalization^138^) on this symmetric contact matrix and plotted splicing RNAs and translation RNAs (columns) versus mRNA exons and introns (rows).

#### Identifying unannotated scaRNAs

We calculated the weighted contact frequency of how often a given RNA contacts scaRNA2. Many of the top hits correspond to Mus musculus (mm10) annotated scaRNAs (e.g. scaRNA9, scaRNA10, scaRNA6, scaRNA7, scaRNA1, scaRNA17, and scaRNA13). Other hits include regions within mRNA introns. We performed BLAST-like Alignment Tool (BLAT, https://genome.ucsc.edu/cgi-bin/hgBlat) on other top hits contacting scaRNA2, including the Trrap intron region and Gon4l1 intron region and found they are homologous to human scaRNA28 and scaRNA26A, respectively. Specifically, the Trrap region in mm10 homologous to scaRNA28 is chr5:144771339-144771531 and the Gon4l region in mm10 homologous to scaRNA26A is chr3:88880319-88880467.

### Analysis of multiway RNA and DNA SPRITE contacts

#### Generating RNA-DNA-DNA Contact Matrices for SPRITE clusters containing an individual or multiple RNAs

To analyze higher-order RNA and DNA contacts in the SPRITE clusters, we generated DNA-DNA contact frequency maps in the presence of specific sets of RNA transcripts. To generate these DNA-DNA contact maps, we first obtained the subset of SPRITE clusters that contained an RNA transcript or multiple transcripts of interest (e.g., nucleolar RNAs, spliceosomal RNAs, scaRNAs satellite RNAs, lncRNA). We then calculated DNA-DNA contact maps for each subset of SPRITE clusters at 100kb and 1Mb resolution by determining the number of clusters in which each pair of genomic bins co-occur. Raw contacts were normalized by SPRITE cluster size by dividing each contact by the total number of reads in the corresponding SPRITE cluster as described above. This resulted in genome-wide DNA-DNA contact frequency maps for each set of RNA transcripts of interest.

#### Aggregate inter-chromosomal maps

For satellite-derived ncRNAs, we also calculated a mean inter-chromosomal contact frequency map. To do this, we converted each 1Mb genomic bin into a percentile bin from 0 to 100 based on its chromosomal position, where the 5’ end is 0 and the 3’ end is 100. We then calculated the contact frequency between all pairs of percentile bins for all pairs of chromosomes. We used these values to calculate a mean inter-chromosomal contact frequency map, which reflects the average contact frequency between each pair of percentile bins between all pairs of chromosomes.

### Satellite-derived ncRNA knockdowns and HP1 measurements

#### LNA transfections

LNA antisense oligonucleotides designed against Major Satellite and Minor Satellite were transfected using Lipofectamine™ RNAiMAX Transfection Reagent according to manufacturer protocol (Thermo Fisher Scientific #13778030). We designed LNAs targeting the forward and reverse strand of the satellite-derived RNAs. These probes, targeting distinct regions of the transcript, were mixed together to a final concentration of 10uM each and 5ul of the mix was transfected to each well of a 24-well plate containing cells. As a control, non-targeting LNA were transfected at the same concentrations. After 48h or 72h in culture, cells were used for further procedures. KD for both LNA were confirmed by RT-qPCRs (**Supplemental Figure 5C-D**).

#### LNA sequences

LNAs were designed by Qiagen. The following sequences were used. Minor Satellite (forward): ACTCACTCATCTAATA, Minor Satellite (reverse): TGGCAAGACAACTGAA, Major Satellite (forward): AGGTCCTTCAGTGTGC, Major Satellite (reverse): ACATTCGTTGGAAACG

#### Reverse transcription and quantitative PCR (RT-qPCR)

Total RNA was extracted from ES cells with Silane beads (Sigma) according to manufacturer conditions and treated with Turbo DNase (Life Technologies) for 15min at 37C to remove genomic DNA. RT reactions were performed according to Superscript II protocol (Thermo Fisher Scientific #18064022) with random 9mer. qPCRs were performed in technical replicates using a Roche Lightcycler and a representative of three biological replicates is shown. Plots were generated using GraphPad software. ddCt values were calculated by normalizing Ct values to GAPDH and to samples transfected with control LNA to compare gene expression differences between samples.

#### qPCR primers used for analysis

GAPDH:CATGGCCTTCCGTGTTCCTA GCCTGCTTCACCACCTTCTT MinS_1: GAACATATTAGATGAGTGAGTTAC GTTCTACAAATCCCGTTTCCAAC MinS_2: GATGGAAAATGATAAAAACC CATCTAATATGTTCTACAGTGTGG MajS_1: GACGACTTGAAAAATGACGAAATC CATATTCCAGGTCCTTCAGTGTGC MajS_2: GCACACTGAAGGACCTGGAATATG GATTTCGTCATTTTTCAAGTCGTC

#### Image analysis of HP1 foci

Image visualization and analysis was performed with Icy software and ImageJ software with a minimum of 10 cells observed per condition. For HP1 foci quantification, we computed a binary mask based on relative intensity threshold (>100 for HP1ß staining replicate 1, >120 for HP1ß replicate 2) in which the relative signal intensity was set from 10 to 200.

### Mapping lncRNA localization

#### Defining lncRNAs

We used Gencode release 95 (GRCm38.p6, https://ftp.ensembl.org/pub/release-95/gtf/mus_musculus/Mus_musculus.GRCm38.95.gtf.gz) to define all lncRNAs in this study. Specifically, we included all annotations with the “lincRNA” or “antisense” biotypes to define all lncRNAs. For example, lncRNAs such as Tsix, Airn, and Kcnq1ot1 are annotated as “antisense” rather than “lincRNA”. We included all lncRNAs that contained coverage in our mouse ES data by filtering the list to those that were contained in at least 10 SPRITE clusters. This yielded a list of 642 lncRNAs.

#### Calculation of chromatin enrichment scores

To determine the extent to which RNA transcripts are in contact with chromatin, we calculated a chromatin enrichment score for each RNA transcript. The chromatin enrichment score is computed as the ratio of the number of SPRITE clusters containing a given RNA that also contains DNA (“chromatin bound”) relative to all SPRITE clusters containing the RNA transcript. We normalize these counts by the SPRITE cluster size in which it was observed. We determined an “expected” DNA to RNA contact ratio by calculating mean DNA to RNA contact ratio across all RNA transcripts. Chromatin enrichment scores were calculated as the natural log of the observed DNA to RNA contact ratio divided by the expected ratio. Positive chromatin enrichment scores indicate RNA transcripts with higher ratios of DNA to RNA contacts than the mean. We performed a similar analysis to calculate enrichment scores for different sets of RNA transcripts. For example, we compute a ribosomal RNA enrichment score based on the ratio of ribosomal RNA contacts to all RNA contacts for a given RNA transcript.

#### lncRNA RNA-DNA genome wide heatmap

We plotted these 642 lncRNAs across the genome at 10Mb resolution. For each lncRNA, we computed the number of SPRITE clusters that co-occur within each 10Mb bin. We then normalized this count by the average contacts across all genomic bins. We refer to this ratio as an enrichment score. This enrichment score is intrinsically normalized for the different expression levels of different lncRNAs. We plotted all bins that have an enrichment value greater than 5-fold. We zoomed in on selected examples and plotted them across the entire genome at 1Mb resolution. In these examples, we plotted the enrichment scores across all values as a continuous bedgraph in IGV.

#### Calculation of lncRNAs enriched around their transcriptional loci

Using these values, we defined a lncRNA as enriched in proximity to its transcriptional locus if it was >20-fold enriched within the 10Mb bin containing its transcriptional loci. At this cutoff, lncRNAs that have very broad distribution patterns across the genome such as Malat1 are excluded, while the vast majority of lncRNAs (596 lncRNAs, 92.8%) are highly enriched around their transcriptional loci.

#### Visualizing proportion of lncRNAs or mRNAs on chromatin

To visually compare the fraction of different RNAs that are retained on chromatin across the genome, we computed a weighted score accounting for the counts within a given genomic bin relative to the total fraction of SPRITE clusters contained off chromatin. Specifically, we identified all SPRITE clusters containing a given RNA and computed the number that also contained a DNA read (on chromatin count) and the number that do not contain DNA (off chromatin count). We computed a score for each genomic bin defined as the number of SPRITE clusters containing an RNA and genomic bin by dividing this count by the total number of SPRITE clusters containing the same RNA that did not have a paired DNA read (off-DNA count). We multiplied this number by 100 to linearly scale values. This score accounts for different abundance levels of different RNAs allowing us to compare them directly to each other and accounts for the proportion of the RNA that is present on chromatin versus off-chromatin.

#### Generating nuclear structure models of lncRNA localization

To visualize the localization of lncRNAs in 3D, we generated 3D models of the genome based on SPRITE DNA-DNA contacts. We modeled each chromosome as a linear polymer composed of N monomers, where N is the number of 1Mb bins on the chromosome. Each chromosome polymer is initialized as a random walk, and then a Brownian dynamics simulation is performed on all chromosomes using an energy function composed of the following forces: 1) a harmonic bond force between adjacent monomers, 2) a spherical confinement force, 3) a repulsive force to prevent monomers from overlapping, 4) an attractive force based on SPRITE contact frequencies to ensure that preferential contacts determined by SPRITE are accurately reflected by the models.

Simulations were performed using the open-source molecular simulation software OpenMM. The outputs of simulations were visualized using Pymol 2 (pymol.org/2). Chromosomes were visualized as cartoon tubes and lncRNAs were visualized by drawing a surface over the genomic regions where lncRNA enrichment was greater than 50-fold over background.

#### FVP treatment and analysis

GRO-seq data from Jonkers *et al*.^139^ were obtained from NCBI GEO (accession GSE48895) and aligned to mm10 using HISAT2. Raw read counts were determined for each gene using deepTools module multiBamSummary for untreated and 50 min FVP conditions. Raw read counts were converted to transcripts per million (TPM) values using a custom Python script, and fold change in TPM was calculated for each gene by dividing 50 min FVP TPM values by untreated TPM values. Cumulative distribution plots were generated using R and box-and-whisker plots were generated using prism.

### Kcnq1ot1 protein binding, perturbations, and gene expression measurements

#### Kcnq1ot1 CRISPR interference

dCas9-4XSID cells were transfected using multiplexed gRNA vector constructs, containing an episomal polyoma origin of replication, puromycin resistance driven by a PGK promoter, and four tandem U6-gRNA cassettes, allowing for simultaneous expression of four sgRNAs. Negative control gRNA sequences recognizing the *Saccharomyces cerevisiae* Upstream Activation Sequence (UAS) and the Tetracycline Response Element (TRE) were multiplexed together (referred to as sgTUUT; gRNAs are as follows: TCTCTATCACTGATAGGGAG, GAGGACAGTACTCCGCTCGG, GCGGAGTACTGTCCTCCGAG, and TCTCTATCACTGATAGGGAG). Four gRNA sequences targeting the Kcnq1ot1 promoter were multiplexed together (referred to as sgKcnq1ot1; gRNAs are as follows: GCCTAGCCGTTGTCGCTAGG, GCCCTGTACTGCATTGAGGT, GCCTGCACAGTAGGATTCCA, and GGAGGATGGGTCGAGTGGCT).

dCas9-4XSID cells were transfected with either sgTUUT or sgKcnq1ot1 and selected for three days with 1μg/ml of puromycin in standard 2i culture conditions. Cells were subsequently passaged and maintained in 0.5μg/ml puromycin for an additional 7 days prior to RNA harvesting. Data presented are from two separate transfections and biological replicates.

#### SHARP binding to Kcnq1ot1 RNA

We transfected an expression vector containing full-length SHARP with an N-terminal Halo-FLAG (HF) fusion protein into mouse ES cells containing a doxycycline inducible Xist gene. Cells were washed once with PBS and then crosslinked on ice using 0.25 J cm−2 (UV2.5k) of UV at 254 nm in a Spectrolinker UV Crosslinker. Cells were then scraped from culture dishes, washed once with PBS, pelleted by centrifugation at 1,500g for 4 min, and flash-frozen in liquid nitrogen for storage at –80°C. We lysed batches of 5 million cells by completely resuspending frozen cell pellets in 1 mL of ice cold iCLIP lysis buffer (50 mM Hepes, pH 7.4, 100 mM NaCl, 1% NP-40, 0.1% SDS, 0.5% Sodium Deoxycholate) supplemented with 1X Protease Inhibitor Cocktail (Promega), 200 U of Murine RNase Inhibitor (New England Biolabs), 20 U Turbo DNase (Ambion), and 1X Manganese/Calcium Mix (0.5mM CaCl2, 2.5 mM MnCl2). Samples were incubated on ice for 10 minutes to allow lysis to proceed. The lysates were then incubated at 37°C for 10 minutes at 1150 rpm shaking on a Thermomixer (Eppendorf). Lysates were cleared by centrifugation at 15,000g for 2 minutes. The supernatant was collected and kept on ice until bound to the HaloLink Resin.

We used 200 μL of 25% HaloLink Resin (50 μL of HaloLink Resin total) per 5 million cells. Resin was washed three times with 2 mL of 1X TBS (50 mM Tris pH 7.5, 150 mM NaCl) and incubated in 1X Blocking Buffer (50 mM HEPES, pH 7.5, 10 μg/mL Random 9-mer, 100 μg/mL BSA) for 20 minutes at room temperature with continuous rotation. After the incubation, resin was washed three times with 1X TBS. The cleared lysate was mixed with 50μl of HaloLink Resin and incubated at 4 °C for 3-16 hrs with continuous rotation. The captured protein bound to resin was washed three times with iCLIP lysis buffer at room temperature and then washed three times at 90°C for 2 minutes while shaking at 1200 rpm with each of the following buffers: 1X ProK/NLS buffer (50 mM HEPES, pH 7.5, 2% NLS, 10 mM EDTA, 0.1% NP-40, 10 mM DTT), High Salt Buffer (50 mM HEPES, pH 7.5, 10 mM EDTA, 0.1% NP-40, 1M NaCl), 8M Urea Buffer (50 mM HEPES, pH 7.5, 10 mM EDTA, 0.1% NP-40, 8 M Urea), and Tween buffer (50 mM HEPES, pH 7.5, 0.1% Tween 20, 10 mM EDTA). Finally, we adjusted the buffer by washing with Elution Buffer (50 mM HEPES, pH 7.5, 0.5 mM EDTA, 0.1% NP-40) three times at 30°C. The resin was resuspended in 83 μL of Elution Buffer and split into a 75 μL (ProK elution) and 8 μL (TEV elution) reaction. 25 μL of 4X ProK/NLS Buffer and 10 μL of ProK were added to the ProK elution tube and the sample was incubated at 50°C for 30 minutes while shaking at 1200 rpm. 2.3 μL of ProTEV Plus Protease (Promega) was added to the TEV Elution and the sample was incubated at 30°C for 30 minutes while shaking at 1200 rpm.

For each experiment, we ensured that we successfully purified the Halo-tagged protein. To do this, the TEV elution sample was mixed with 1X LDS Sample Buffer (Invitrogen) and 1X Reducing Agent (Invitrogen) and heated for 6 minutes at 70°C. The sample was run on a 3-8% Tris Acetate Gel (Invitrogen) for 1 hour at 150 V. The gel was transferred to a nitrocellulose membrane using an iBlot Transfer Device (Invitrogen). The nitrocellulose membrane was blocked with Odyssey Blocking Buffer (LI-COR) for 30 minutes. We incubated the membrane in Anti-FLAG mouse monoclonal Antibody (Sigma, F3166) and V5 rabbit polyclonal antibody (Santa Cruz, sc-83849-R) at a 1:2500 dilution for 2 hours at room temperature to detect the protein. We visualized the protein by incubating the membrane in 1:17,500 dilution of both IRDye 800CW Goat anti-Rabbit IgG (LI-COR, 925-32210) and IRDYE 680DR Goat anti-Mouse IgG (LI-COR, 925-68070) for 1 hour at room temperature followed by imaging on a LICOR Odyssey.

RNA was purified from the Proteinase K elution sample and an RNA-Seq library was constructed as previously described. Briefly, after proteinase K elution, the RNA was dephosphorylated (Fast AP) and cyclic phosphates removed (T4 PNK) and then cleaned up on Silane beads as previously described. The RNA was then ligated to an RNA adapter containing a RT primer binding site. The ligated RNA was reverse transcribed (RT) into cDNA, the RNA was degraded using NaOH, and a second adapter was ligated to the single stranded cDNA. The DNA was amplified and Illumina sequencing adaptors were added by PCR using primers that are complementary to the 3’ and 5’ adapters. The molarity of PCR amplified libraries was measured by Agilent Tapestation High Sensitivity DNA screentapes and all samples were pooled at equal molarity. The pool was then purified and size selected on a 2% agarose gel and cut between 150-700 nts. The final libraries were measured by Agilent Bioanalyzer and Qubit high sensitivity DNA to determine the loading density of the final pooled sample. Pooled samples were paired-end sequenced on an Illumina HiSeq 2500 with read length 35 x 35nts.

Sequencing reads were trimmed to remove adaptor sequences and any bases containing a quality scores <10 using Trimmomatic^140^. We filtered out all read-pairs where either read was trimmed to <25 nucleotides. We excluded PCR duplicates using the FastUniq tool^141^. The remaining reads were then aligned to Ribosomal RNAs (rRNAs) using the Tagdust program^142^ with a database of 18S, 28S, 45S, 5S, 5.8S sequences. TagDust was chosen because it allowed more permissive alignments to rRNA reads that contained mismatches and indels due to RT errors induced by rRNA post-transcriptional modifications. The remaining reads were then aligned to the mouse genome using STAR aligner^143^. Only reads that mapped uniquely in the genome were kept for further analysis.

#### Genetic deletion of SHARP Binding Site in Kcnq1ot1

F1 2-1 line were CRISPR-targeted with gRNAs targeting the SHARP-Binding Site (SBS) (SHARP Binding Site Coordinates: mm10 - chr7:143,295,789-143,296,455; gRNA sequences were ATGCACCATCATAGACCACG and TCATAGCCTCCCCCTCCTCG). Following selection using 1μg/ml of puromycin in standard 2i culture conditions, transfected cells were allowed to recover in standard 2i media prior to sub-cloning. Clone were subsequently screened using genomic DNA PCR, using primers flanking the deletion region (CAGCATCTGTCCAATCAACAG and GCAAAATACGAGAACTGAGCC respectively). In contrast to the wild type 1048bp band, successfully targeted alleles would produce 305bp band. Sub-clones homozygous for the targeted allele were subject to RT-qPCR and GAPDH-normalized gene expression was further normalized to the F1 parent line).

#### HDAC inhibitor treatment

The inducible Kcnq1ot1 cell line were treated with either DMSO (control) or 5μM TSA in fresh 2i media or 2μg/ml doxycycline in standard 2i. RNA was extracted, reverse transcribed, and qPCR was performed. CT values were normalized to GAPDH to compare gene expression differences between induced and non-induced samples within the same pharmacologic condition (i.e. GAPDH-normalized “Induced DMSO” to GAPDH-normalized “Non-Induced DMSO Vehicle) to generate fold gene expression ratios. RT-qPCR data presented is summarized from two separate replicate experiments

